# Geometric Sketching Compactly Summarizes the Single-Cell Transcriptomic Landscape

**DOI:** 10.1101/536730

**Authors:** Brian Hie, Hyunghoon Cho, Benjamin DeMeo, Bryan Bryson, Bonnie Berger

## Abstract

Large-scale single-cell RNA-sequencing (scRNA-seq) studies that profile hundreds of thousands of cells are becoming increasingly common, overwhelming existing analysis pipelines. Here, we describe how to enhance and accelerate single-cell data analysis by summarizing the transcriptomic heterogeneity within a data set using a small subset of cells, which we refer to as a geometric sketch. Our sketches provide more comprehensive visualization of transcriptional diversity, capture rare cell types with high sensitivity, and accurately reveal biological cell types via clustering. Our sketch of umbilical cord blood cells uncovers a rare subpopulation of inflammatory macrophages, which we experimentally validated *in vitro*. The construction of our sketches is extremely fast, which enabled us to accelerate other crucial resource-intensive tasks such as scRNA-seq data integration. We anticipate that our algorithm will become an increasingly essential step when sharing and analyzing the rapidly-growing volume of scRNA-seq data and help enable the democratization of single-cell omics.

## INTRODUCTION

Improvements in the throughput of single-cell profiling experiments, especially droplet-based single-cell RNA-sequencing (scRNA-seq), have resulted in data sets containing hundreds of thousands of cells (Angerer et al., 2017; Macosko et al., 2015; Zheng et al., 2017), with hundreds to thousands of gene expression measurements per cell. As these sequencing pipelines become cheaper and more streamlined, experiments profiling tens of millions of cells may become ubiquitous in the near future (Angerer et al., 2017), and consortium-based efforts like the Human Cell Atlas plan to profile billions of cells (Rozenblatt-Rosen et al., 2017). Leveraging this data to improve our understanding of biology and disease will require merging and integrating many cells across diseases and tissues (Hie et al., 2018), resulting in reference data sets with massive numbers of cells. Unfortunately, the sheer volume of scRNA-seq data being generated is quickly overwhelming existing analytic procedures, requiring prohibitive runtime or memory usage to produce meaningful insights (Angerer et al., 2017; Butler et al., 2018; Lun et al., 2016a). This bottleneck limits the utility of these emerging large data sets to researchers with access to expensive computational infrastructure, and makes quick exploratory analyses impossible even for these researchers.

Here, we present a novel approach to facilitate the sharing and analysis of large-scale single-cell data sets by intelligently selecting a small subset of data (referred to as a “sketch”) that comprehensively represents the transcriptional heterogeneity within the full data set. Because of their vastly reduced computational overhead, our sketches can be efficiently shared among researchers and be used to quickly identify important patterns in the full data set to be followed up with in-depth analyses.

Currently, researchers often uniformly downsample a data set to obtain a small subset to accelerate the initial data analysis (10x Genomics, 2017). Although this simple approach could be used to generate sketches of single-cell data sets, it is highly prone to removing rare cell types and negates the advantage of performing large-scale scRNA-seq experiments in the first place. Alternative sampling approaches that better consider the structure of the data, including *k*-means++ sampling (Arthur and Vassilvitskii, 2007) and spatial random sampling (SRS) (Rahmani and Atia, 2017a), have not yet been applied to the problem of obtaining informative sketches of scRNA-seq data to our knowledge. However, these data-dependent sampling techniques not only lack the ability to efficiently scale to large data sets, but also lack robustness to different experimental settings and produce highly unbalanced sketches that are ill-suited for downstream scRNA-seq analyses as we demonstrate in our experiments.

The key insight behind our novel sampling approach is that common cell types form dense clusters in the gene expression space, while rarer subpopulations may still inhabit comparably large regions of the space but with much greater sparsity. Rather than sample cells uniformly at random, we sample *evenly across the transcriptomic space*, which naturally removes redundant information within the most common cell types and preserves rare transcriptomic structure contained in the original data set. We refer to our sampling method as “geometric sketching” because it obtains random samples based on the geometry, rather than the density, of the data set (**Figure 1**).

**Figure 1.**
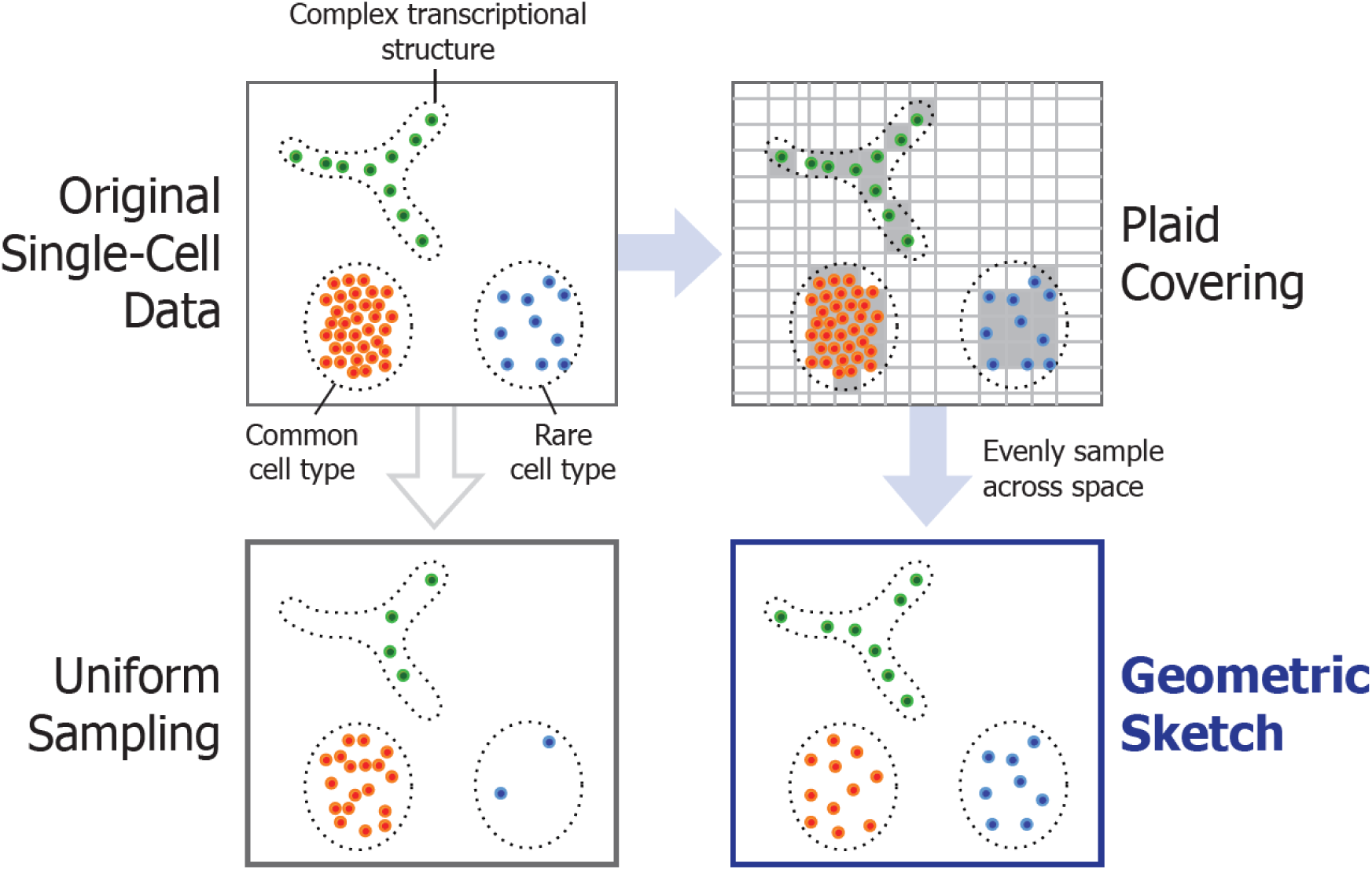
Illustration of Geometric Sketching. We first cover the data points with equal-sized boxes (which we refer to as a *plaid covering*) to approximate their geometry, then sample data points by first spreading the desired total sample count over the boxes as evenly as possible, then choosing the assigned number of samples within each box uniformly at random. The resulting sketch more evenly covers the landscape of the data compared to uniform sampling of points, where the latter is more prone to omitting rare cell types or transcriptional patterns.

Geometric sketching is extremely efficient, sampling from data sets with millions of cells in a matter of minutes and with an asymptotic runtime that is close to linear in the size of the data set. We empirically demonstrate that our algorithm produces sketches that more evenly represent the transcriptional space covered by the data. We further show that our sketches enhance and accelerate downstream analyses by preserving rare cell types, producing visualizations that broadly capture transcriptomic heterogeneity, facilitating the identification of cell types via clustering, and accelerating integration of large scRNA-seq data sets. Moreover, we demonstrate how the sensitivity of geometric sketching to rare transcriptional states allows us to identify a previously unknown rare subpopulation of inflammatory macrophages in a human umbilical cord blood data set, providing insight into a fundamental immunological process. As the size of single-cell data grows, geometric sketching will become increasingly crucial for the democratization of large-scale single-cell experiments, making key analyses tractable even for researchers without expensive computational resources.

## RESULTS

### Overview of Our Geometric Sketching Algorithm

The overall approach taken by the geometric sketching algorithm is illustrated in **Figure 1**. Geometric sketching aims to select a subset of cells (i.e., a sketch) from a large scRNA-seq data set such that the subset accurately reflects the full transcriptomic heterogeneity, where the small size of the sketch enables fast downstream analysis. In order to effectively summarize the diversity of gene expression profiles within a data set, the first step of our algorithm is to approximate the geometry of the transcriptomic space inhabited by the input data as a union of fixed shapes that admit succinct representation. In our work, we approximate the data with a collection of equal-sized, non-overlapping, axis-aligned boxes (hypercubes), which we refer to as a *plaid covering*. We use boxes instead of spheres to obtain a highly efficient greedy covering algorithm that helps us better cope with the increasing volume of scRNA-seq data. Once the geometry of data is approximated via plaid covering, we sample cells by first spreading the desired total sample count over the covering boxes as evenly as possible (based on a random ordering of the boxes), then choosing the assigned number of samples within each box uniformly at random. This process allows the samples to more evenly cover the gene expression landscape of the data, naturally diminishing the influence of densely populated regions and increasing the representation of rare transcriptional states. A more detailed description and theoretical analysis of our approach is provided in **Methods**.

### Geometric Sketches Evenly Summarize the Transcriptomic Landscape

We first sought to quantify how well geometric sketching is able to evenly represent the original transcriptomic space by measuring the Hausdorff distance from the full data set to a geometric sketch (**Methods).** Intuitively, a low Hausdorff distance indicates that the points in a sketch are close to all points in the remainder of the data set within the transcriptomic space, while a high Hausdorff distance indicates that there are some cells in the full data set that are not well represented within the sketch. We benchmarked geometric sketching against uniform sampling as well as more complex, data-dependent strategies: *k*-means++ sampling (Arthur and Vassilvitskii, 2007) and spatial random sampling (SRS) (Rahmani and Atia, 2017a). Note that, to our knowledge, neither of these non-uniform sampling approaches have been previously considered for the problem of downsampling single-cell data sets. *k*-means++ works by randomly choosing an initial sample, then repeatedly sampling the next point such that more distant points from the current sample set have higher probability. SRS works by projecting the data onto the unit ball, sampling points uniformly across the surface of the ball, and picking the closest example from the data set to each of those random points.

We used four scRNA-seq data sets of varying sizes and complexities to assess our method (**Methods; Supplementary Table 1-4**): a 293T/Jurkat mixture with 4,185 cells (Zheng et al., 2017); a PBMC data set with 68,579 cells (Zheng et al., 2017); a developing and adolescent mouse central nervous system (CNS) data set with 465,281 cells (Zeisel et al., 2018); and an adult mouse brain data set with 665,858 cells (Saunders et al., 2018). In all cases, we observed that geometric sketching obtains substantially better improvement under a robust Hausdorff distance measure (**Methods**) than uniform sampling and the other data-dependent sampling methods, SRS and *k*-means++ (**Figure 2**). The improvement in Hausdorff distance was consistent across sketch sizes ranging from 2% to 10% of the full data set, providing quantitative evidence that our algorithm more evenly samples over the geometry of the data set than do other methods.

**Figure 2.**
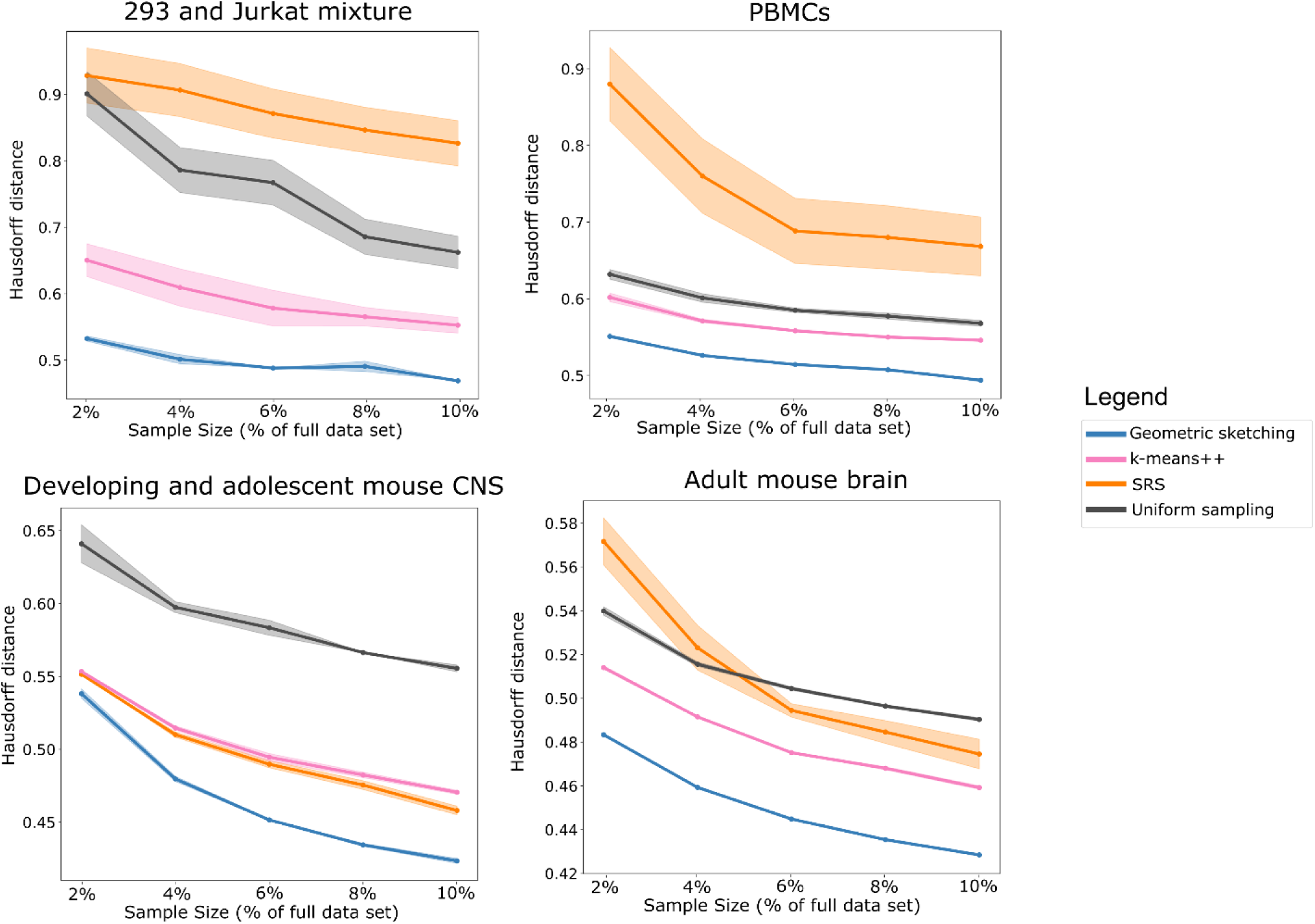
Geometric Sketching Yields More Even Coverage of the Transcriptomic Space. In our experiments, the Hausdorff distance measures the maximum distance from any point in the data set to its closest point in the sketch; a lower Hausdorff distance indicates that the points represented by a sketch are in general closer to all of the points in the remainder of the data set. Geometric sketching results in consistently lower Hausdorff distances than other sampling methods across a large number of sketch sizes and data sets. We use a robust Hausdorff distance that is less sensitive to small numbers of outlier observations (**Methods**). Solid lines indicate means and shaded areas indicate standard error across 10 random trials for geometric sketching and uniform sampling and 4 random trials for *k-*means++ and SRS (due to long runtimes).

### Visualization of Geometric Sketches Reveals Transcriptional Diversity

We next set out to assess the ability of our geometric sampling approach to improve the low-dimensional visualization of scRNA-seq data, a common exploratory (and often computationally expensive) initial step in single-cell genomic analysis. From our two largest data sets of mouse nervous system, containing 465,281 and 665,858 cells each, we used a 2-dimensional *t*-SNE (Van Der Maaten and Hinton, 2008) to visualize a sketch containing 2% of the total data set (sampled without replacement) obtained by geometric sketching.

The results, shown in **Figure 3**, illustrate that the relative representations of cell types in geometric sketches can have striking differences compared to uniformly downsampled data sets. For instance, when obtaining a sketch of 2% of the data set of adult mouse neurons (Saunders et al., 2018), clusters of macrophages, endothelial tip cells, and mural cells have only 59, 117, and 336 cells, respectively, in the uniform sample out of 1695, 3818, and 12083 cells in the full data, respectively. In contrast, these cell types have 326, 1022, and 875 cells, respectively, in the geometric sketch of the same size. Although these cell types are rare compared to neurons (428,051 cells in the full data set), their substantially increased representation in our sketch suggests they inhabit a comparatively large portion of the transcriptional space. Similarly, on a data set of 465,281 cells from the developing and adolescent mouse central nervous system (CNS) (Zeisel et al., 2018), we also observed a more balanced composition of cell types as determined by the original study’s authors (**Figure 3**). The rarest cell types are also more consistently represented in a geometric sketch than in sketches obtained by SRS or *k*-means++ (**Supplementary Fig. 1A; Supplementary Table 5-6**). We also visualize the data using uniform manifold approximation and project (UMAP), an alternative method for computing 2-dimensional visualization embeddings (McInnes and Healy, 2018) with similar results as those produced by our *t*-SNE experiments (**Supplementary Fig. 1B**).

**Figure 3.**
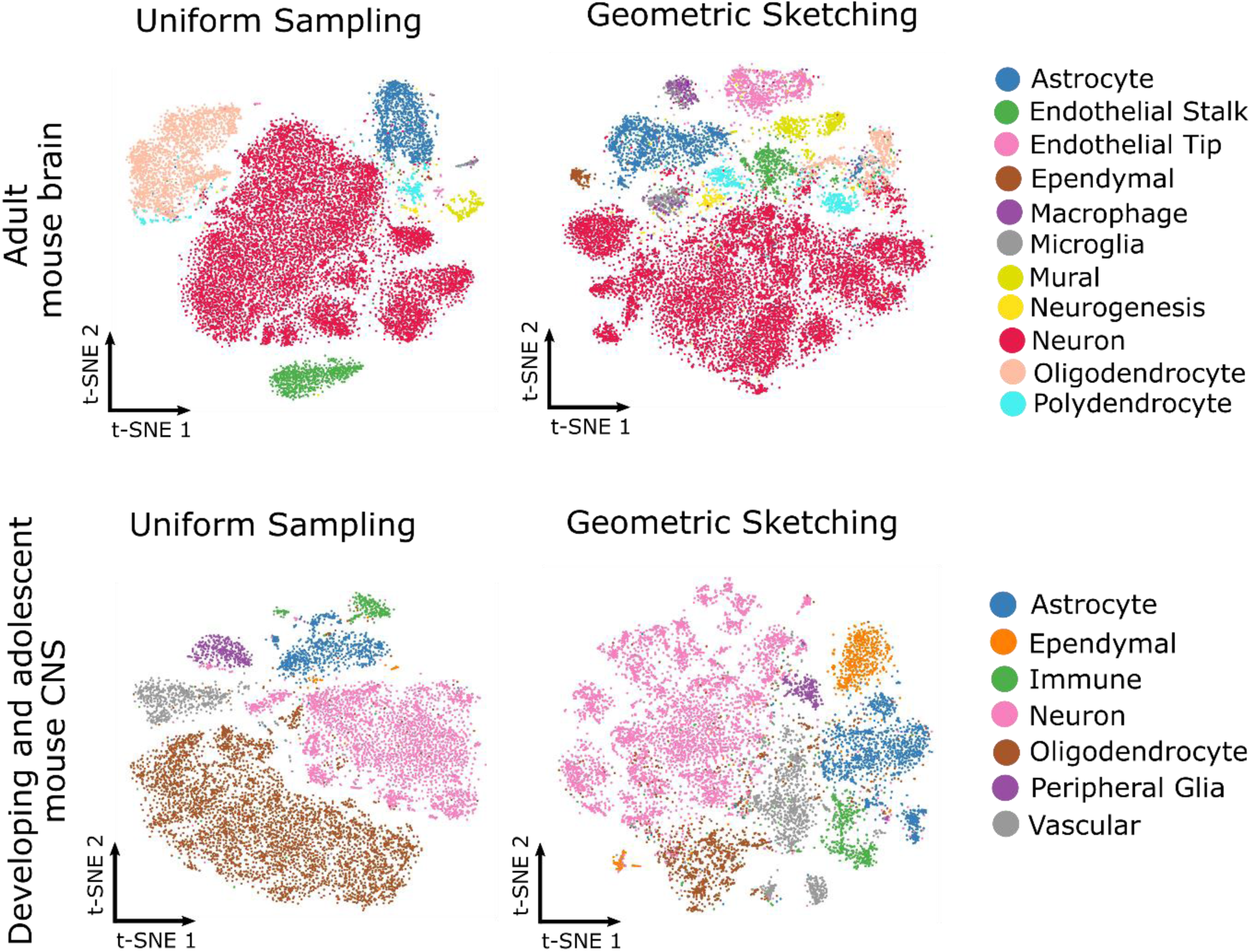
Geometric Sketching Produces More Balanced Summaries of Transcriptional Landscape. *t*-SNE visualizations of sketches containing 2% of the cells from the adult mouse brain (Saunders et al., 2018) and from the developing and adolescent mouse CNS (Zeisel et al., 2018) using uniform random sampling and geometric sketching, with increased representation of rare cell types in the geometric sketch. Numbers of cells from each cell type are given in **Supplementary Tables 5-6**. Uniform sampling, which does not evenly consider the transcriptional space, produces visualizations that are poor at capturing transcriptional heterogeneity. Geometric sketching substantially underrepresents oligodendrocytes in both data sets compared to uniform sampling, which is expected given the low transcriptional heterogeneity among oligodendrocytes as quantified by differential entropy (**Methods; Supplementary Tables 3-4**). Visualizations based on other sampling approaches as well as a different visualization method are provided in **Supplementary Fig. 1**.

We note that our sampling algorithm is completely unsupervised and has no knowledge of the cell type labels, but naturally obtains a balanced composition of cell types by sampling more evenly across the entire transcriptional space. Indeed, on artificial data in which we controlled the relative volumes and densities of the clusters, geometric sketching samples the clusters proportionally to their relative volumes rather than their frequencies in the full data set (**Supplementary Fig. 2**), suggesting that the composition of different cell types in a geometric sketch more closely reflects the transcriptional variability of individual clusters rather than their frequency in the overall population. Our visualizations therefore reflect a geometric “map” of the transcriptional variability within a data set, allowing researchers to more easily gain insight into rarer transcriptional states.

### Rare Cell Types Are Better Preserved Within Geometric Sketches

As suggested by the above results, one of the key advantages of our algorithm is that it naturally increases the representation of rare cell types with sufficient transcriptomic heterogeneity in the subsampled data. Using the four data sets mentioned above, which include cell type labels provided by the original study authors, we evaluated the ability of our method to preserve the rarest cell type within each data set. In particular, we focused on 28 293T cells (0.66% of the total number of cells in the data set) in the 293T/Jurkat mixture, 262 dendritic cells (0.38%) in the PBMC data set, 1695 macrophages (0.25%) among the adult mouse brain cells, and 2777 ependymal cells (0.60%) among the mouse CNS cells. In all data sets, the rare cell types are substantially more represented in the sketch obtained by our algorithm compared to other sampling techniques (**Figure 4**). For example, a sketch that is 2% the size of the 665,858 mouse brain cells contains an average of 281 macrophages compared to only 31 cells from uniform sampling. Geometric sketching is able to better preserve rare cell types because the extent of transcriptional variation among rare cells is similar to that of common cells. To this end, we used the differential entropy of a multivariate Gaussian fit to each cell type as a proxy to its transcriptional diversity (**Methods**; **Supplementary Table 1-4**). We also note that, within the geometric sketch, almost all of the rare cell types in each data set have increased representation compared to the full data, where the representation of rare cell types gradually converges to that of uniform sampling as the sketch size increases (**Supplementary Fig. 3**).

**Figure 4.**
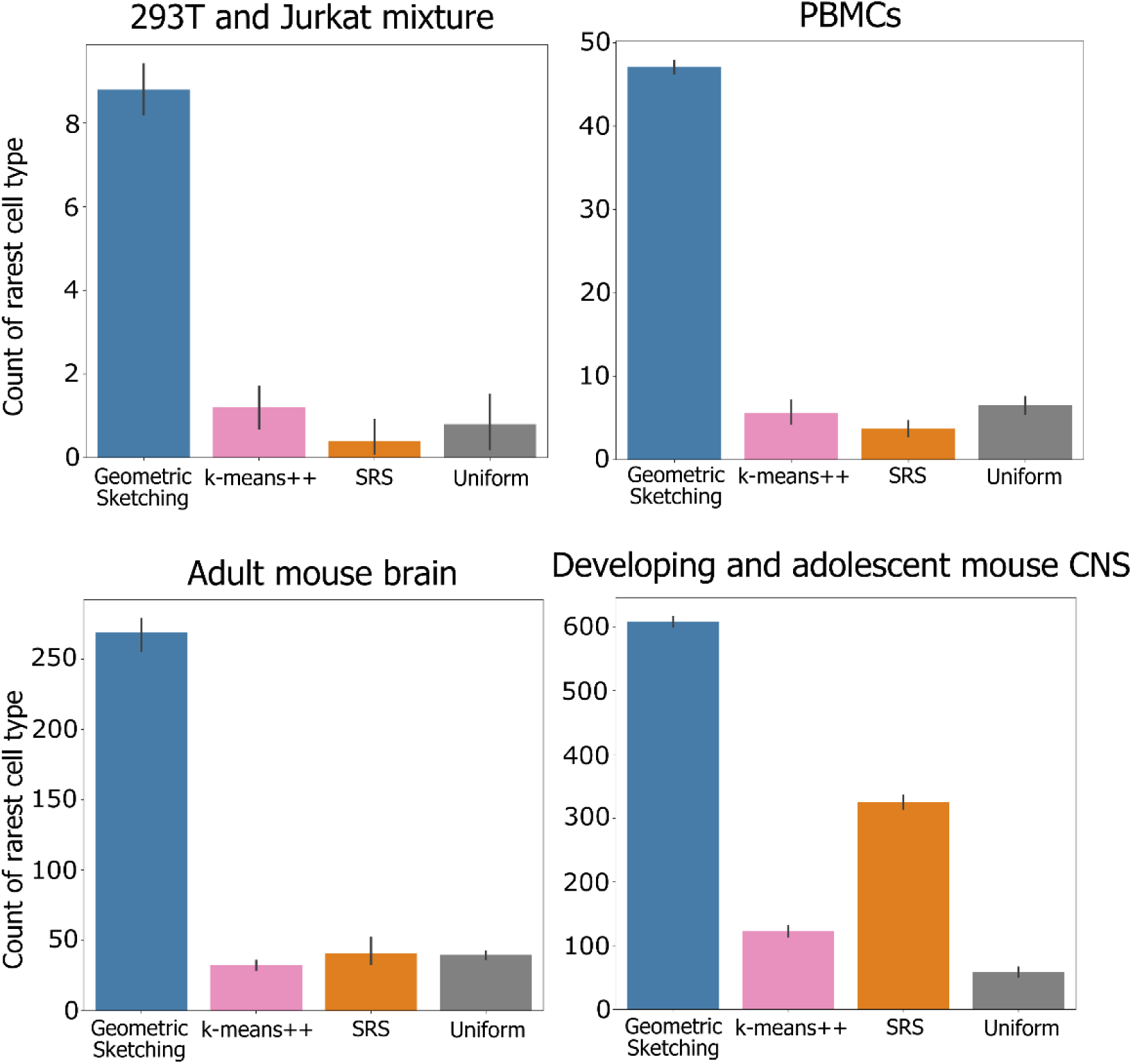
Geometric Sketching Preserves Rare Cell Types in the Subsampled Data. In sketches containing 2% of the total data set, we counted the number of cells that belong to the rarest cell type in each data set: 293T cells (0.66% of total cells) in a 293T/Jurkat mixture, dendritic cells (0.38% of total) in a data set of 68k PBMCs, macrophages (0.25% of total) in a data set of adult mouse brain cells, and ependymal cells (0.60% of total) in a data set of developing and adolescent mouse CNS cells. Higher count indicates increased representation of the rare cell type in the sketch. Bar height indicates means and error bars indicate standard error across 10 random trials for geometric sketching and uniform sampling and 4 random trials for *k-* means++ and SRS (due to long runtimes). Comparison of rare cell type representation over different sketch sizes is shown in **Supplementary Fig. 3**.

### Clustering of Geometric Sketches Better Recapitulates Biological Cell Types

Since the samples produced by our algorithm consist of a more balanced composition of cell types, including rare cell types, we also reasoned that clustering analyses should be able to better distinguish these cell types within a geometric sketch compared to uniform downsampling. To assess this capability, we first clustered the sketches using the standard graph-based Louvain clustering algorithm (Blondel et al., 2008). Then, we transferred cluster labels to the rest of the data set via *k*-nearest-neighbor classification and assessed the agreement between our unsupervised cluster labels and the biological cell type labels provided by the original studies (**Methods**). We quantified the clustering accuracy via balanced adjusted mutual information (BAMI), our newly proposed metric for evaluating clustering quality when the ground truth clusters are highly imbalanced, which is often the case for scRNA-seq data sets. BAMI balances the terms in adjusted mutual information (Vinh et al., 2010) to equally weight each of the ground truth clusters, preventing rare cell types from having only negligible contribution to the performance metric. We also provide results for adjusted mutual information, without our balancing technique, which are largely consistent with our comparisons based on BAMI (**Supplementary Fig. 4**).

On a variety of real scRNA-seq data sets, our algorithm’s sketches recapitulate the biological cell types consistently better than uniform sampling (**Figure 5**). Although two other data-dependent sampling methods, SRS and *k*-means++, achieve performance comparable to our method in a few cases, only geometric sketching obtains competitive performance across all data sets, suggesting that our method is reasonably robust to different experimental settings. Notably, because our sketches are drawn without replacement, clustering scores can become closer to those of uniform samples as the size of the sketch increases; this may explain the diminishing performance of our method with increasing sketch size on the mixture of 293T cells and Jurkat cells (**Figure 5**). Still, we note our substantial advantage even on this data set using very small sketches that select as low as 2% of the full data set. Moreover, the overall improvement in clustering consistency could become more pronounced as more fine-grain clusters become available as ground truth in light of the enhanced representation of rare transcriptional states within geometric sketches.

**Figure 5.**
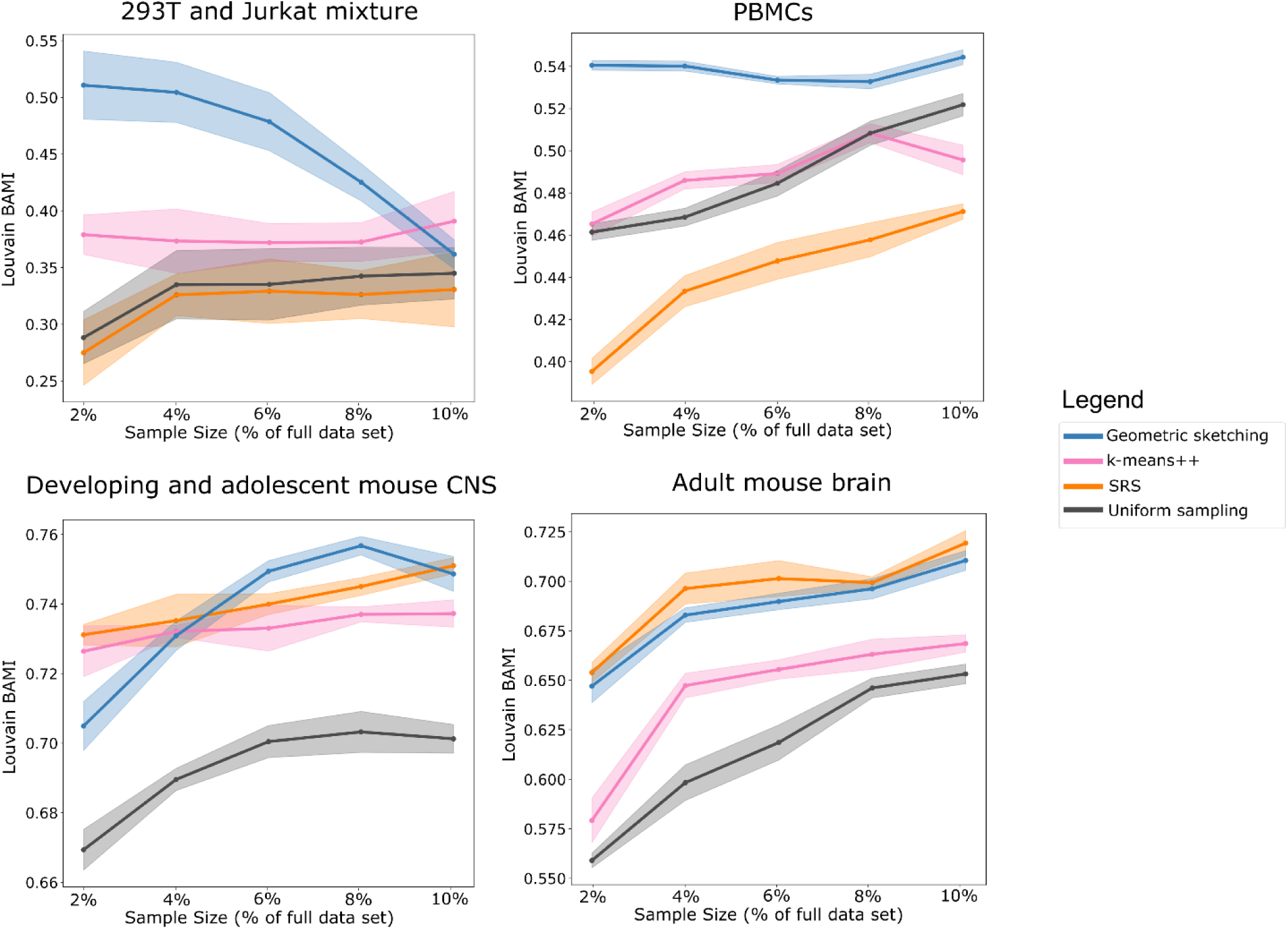
Geometric Sketching is Consistently Effective at Distinguishing Biological Cell Types via Clustering. Louvain clustering was applied to a subsample of the data set obtained by geometric sketching or other baseline algorithms. We transferred the cluster labels to the full data set using a *k*-nearest-neighbor classifier fit to the sketch, then measured the balanced adjusted mutual information (BAMI) between the unsupervised cluster labels and the labels corresponding to biological clusters provided by each previous study (**Methods**). Higher score indicates greater agreement between unsupervised clustering and biological cell type labels. Solid lines indicate means and shaded areas indicate standard error across 10 random trials for geometric sketching and uniform sampling and 4 random trials for *k-*means++ and SRS (due to long runtimes). Unsupervised clustering of geometric sketches consistently recapitulates biological cell types better than clustering results obtained by uniform sampling. Other non-uniform sampling methods, *k*-means++ and SRS, show performance comparable to ours in a few cases, but only geometric sketching obtains competitive performance across all settings. Note that only uniform sampling has been previously considered for the problem of subsampling single-cell data. Because samples are drawn without replacement, clustering accuracy may approach that of uniform sampling as the sketch size increases, as is the case in the 293T/Jurkat mixture experiments.

### Geometric Sketching Assists in the Discovery of a Rare Population of Inflammatory Macrophages

Because geometric sketching of large data sets highlights rare transcriptional states, certain subpopulations of cells that are difficult to identify when analyzing the full data set may become discoverable within a geometric sketch. To test this in practice, we analyzed a data set of 254,941 cells taken from human umbilical cord blood without cell type labels (**Methods**). We computed a geometric sketch of 20,000 cells and clustered the sketch via the Louvain community detection algorithm. Among the putative macrophage clusters with elevated expression of macrophage-specific marker genes, including *CD14* and *CD68* (Khazen et al., 2005), we found a comparatively rare cluster of macrophages defined by the marker genes *CD74, HLA-DRA, B2M*, and *JUNB* (AUROC > 0.90; **Methods**) (**Figure 6**). We hypothesized that this cluster corresponds to *inflammatory* macrophages, since each of its marker genes has been implicated in macrophage activation in response to inflammatory stimuli: *CD74* encodes the receptor for macrophage migration inhibitory factor (MIF) (Leng et al., 2003), a pro-inflammatory signal (Morand et al., 2006; Santos and Morand, 2009); HLA-DR has elevated expression in classically pro-inflammatory M1-macrophages (Helm et al., 2014); increased *B2M* has been demonstrated in murine bone marrow derived macrophages after LPS stimulation (Tanaka et al., 2017); and *JUNB* has been implicated as a key transcriptional modulator of macrophage activation (Fontana et al., 2015) and is upregulated by MIF (Calandra and Roger, 2003). We did not observe major differences in the number of unique genes between this rare cluster and the rest of the macrophages (**Supplementary Fig. 5**), so these differences in gene expression are most likely not an artifact of variable data sparsity or dropout.

**Figure 6.**
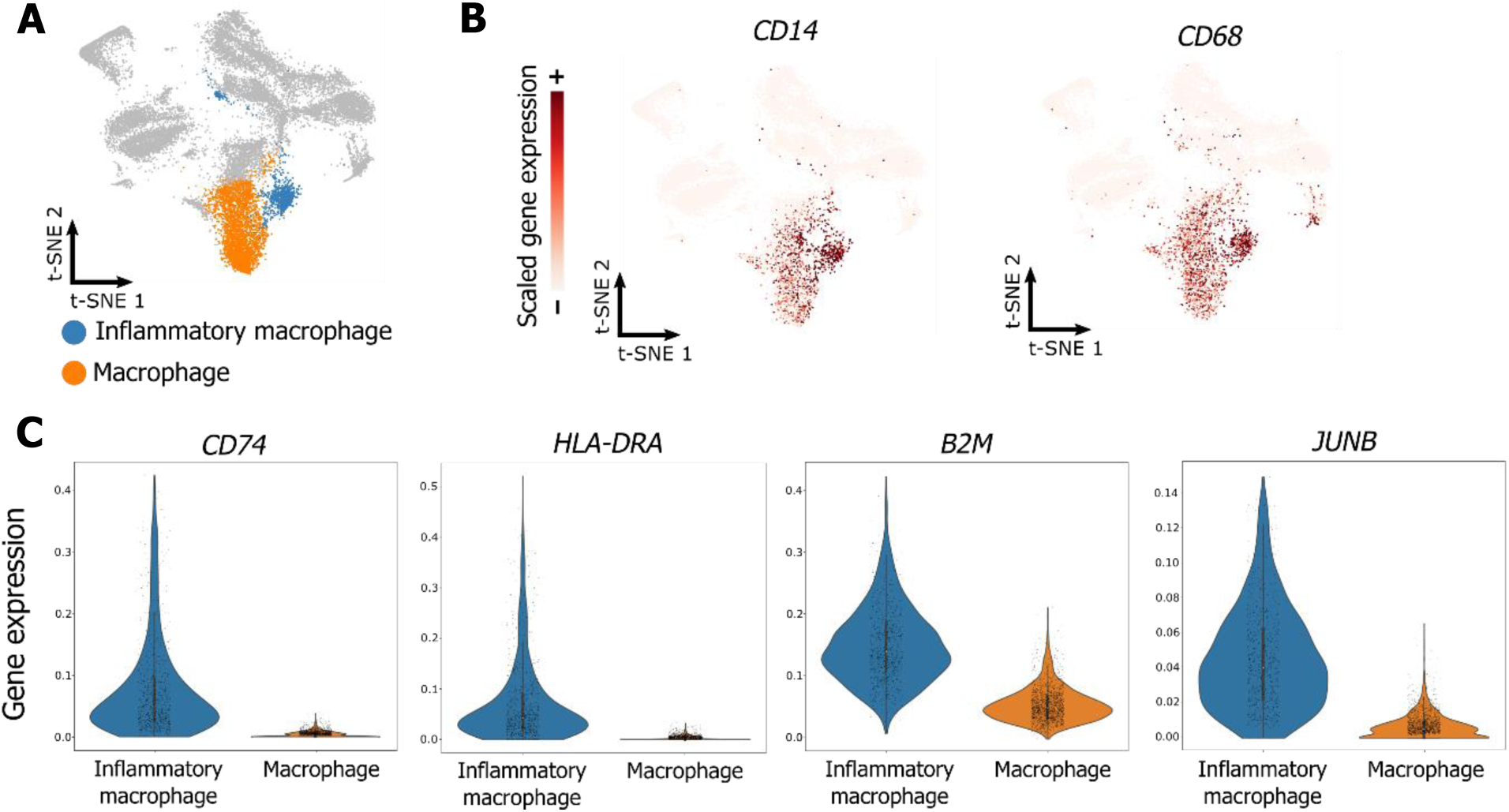
Novel Subpopulation of Inflammatory Macrophages Identified Using Geometric Sketching. A geometric sketch of 20,000 cells was obtained from a full data set of 254,941 cells from human umbilical cord blood. Analysis of clusters obtained by the Louvain community detection algorithm reveals multiple clusters of macrophages (**A**), defined by *CD14* and *CD68* marker gene expression (**B**). A rare subpopulation of these macrophages is in turn defined by inflammatory marker gene expression (*CD74, HLA-DRA, B2M*, and *JUNB*) (**C**), providing insight into an important but comparatively rarer immunological process.

We sought further confirmation of this rare expression signature in macrophages by conducting a separate scRNA-seq study of an *in vitro* model of macrophage inflammation in which human CD14+ monocytes were polarized with GM-CSF to induce an inflammatory response (**Methods**). We compared this data to a similar scRNA-seq data set of macrophages but with M-CSF stimulation (Hie et al., 2018) to induce an anti-inflammatory polarization. Expression of all four marker genes we identified (*CD74, HLA-DRA, B2M*, and *JUNB*) was significantly elevated in GM-CSF-derived (*n* = 354 cells) macrophages compared to the M-CSF-derived (*n* = 1107 cells) macrophages (one-sided Welch’s *t*-test *P* = 4e-34 for *CD74, P* = 1e-29 for *HLA-DRA, P =* 3e-46 for *B2M*, and *P* = 1e-13 for *JUNB*), increasing our confidence in these marker genes as a signature of inflammation. Additional *in vivo* confirmation of these markers, along with more in-depth study of macrophage subpopulations, will help reveal insight into inflammation and ways to modulate inflammatory processes in response to disease.

When we applied the same clustering procedure to either the full data set or a uniform subsample, the clustering algorithm did not assign a separate cluster to inflammatory macrophages but rather placed all macrophages into a single cluster, likely because of the relative scarcity of this cell type compared to the large cluster of inactive macrophages. These results provide additional evidence that geometric sketches contain a richer variety of transcriptional states and can therefore assist researchers in identifying interesting but rare biological structure.

### Geometric Sketching Has Significantly Better Scalability to Large Data Sets Than Other Sophisticated Sampling Strategies

Not only does geometric sketching lead to more informative sketches of the single-cell data, it is also dramatically faster than other non-uniform sampling methods, which is imperative since researchers stand to gain the most from sketches of very large data sets. Geometric sketching has an asymptotic runtime that is close to linear in the size of the data set (**Methods**) and, when benchmarked on real data sets, is more than an order of magnitude faster than non-uniform methods and has a negligible dependence on the number of samples specified by the user, unlike *k*-means++ and SRS (**Figure 7**). On our largest data set of 665,858 cells, our sampling algorithm takes an average of around 5 minutes (**Figure 7**); in contrast, *k*-means++ takes 3 hours and spatial random sampling (SRS) takes 5.5 hours when subsampling 10% of the cells. On a simulated benchmark data set of 10 million data points (**Methods**), geometric sketching subsamples 20,000 cells after an average time of 67 minutes, demonstrating practical scalability to data sets with hundreds of millions of cells (**Figure 7**). Although uniform sampling is trivially the most efficient technique since it does not consider any properties of the underlying data set, our algorithm is both efficient and produces high quality samples that more accurately represent the underlying transcriptomic space as we demonstrated above. Notably, our runtime comparison does not include the standard preprocessing step of (randomized) principal component analysis (PCA), which we uniformly applied to all methods and whose runtime as well as scalability are comparable to our geometric sketching step (**Methods**; **Supplementary Fig. 8**).

**Figure 7.**
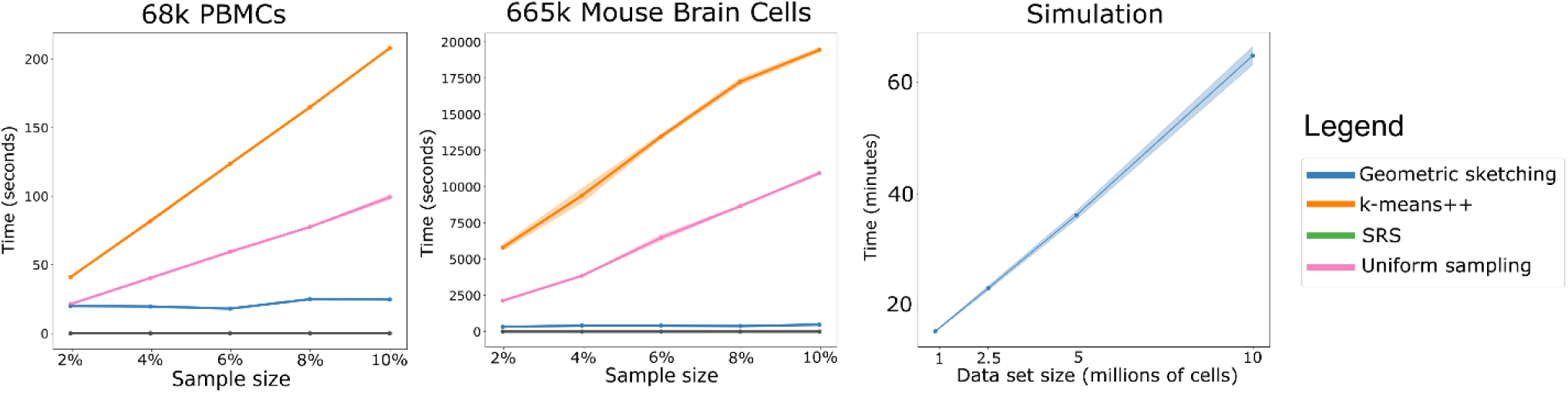
Geometric Sketching Efficiently Scales to Large Single-Cell Data Sets. Geometric sketching is substantially more efficient than other data-dependent subsampling approaches, SRS and *k*-means++. Although uniform sampling is fastest because it does not consider any properties of the data set, geometric sampling obtains a sketch that preserves transcriptional heterogeneity while running in close to linear time in the size of the data, largely independent of the requested number of samples. Solid lines indicate means and shaded areas indicate standard error across 10 random trials for geometric sketching and uniform sampling and 4 random trials for *k-*means++ and SRS (due to long runtimes). Geometric sketching has a practical runtime of around 67 minutes when sampling 20,000 cells from a simulated data set with 10 million cells, which was obtained by resampling from a data set of mouse CNS cells (Zeisel et al., 2018).

**Figure 8.**
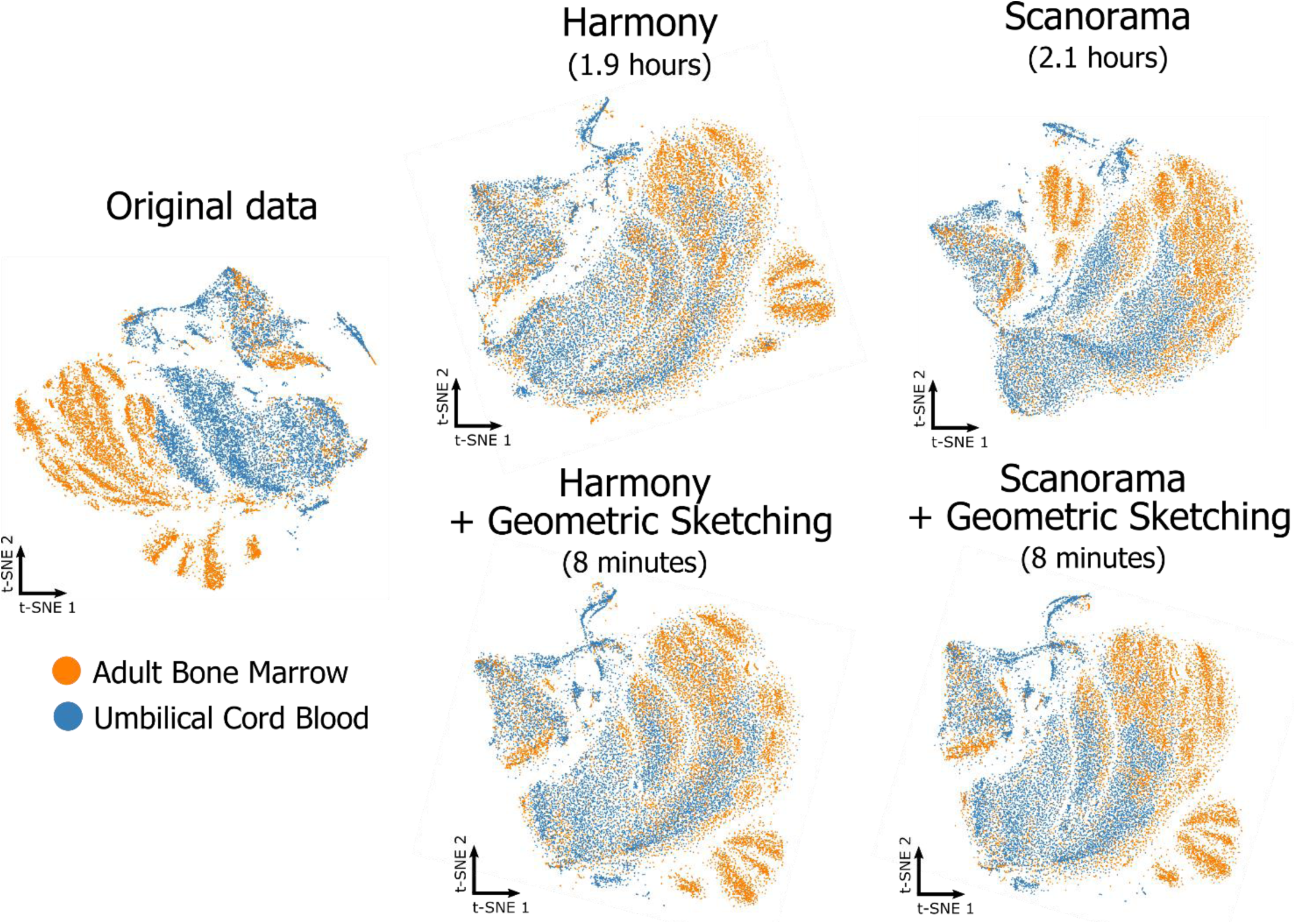
Geometric Sketching Accelerates Single-Cell Data Integration Tools. Geometric sketching can help accelerate existing tools for scRNA-seq data integration. We use two existing algorithms for scRNA-seq integration, namely Harmony (Korsunsky et al., 2018) and Scanorama (Hie et al., 2018), but note that our approach works for other integrative algorithms as well. Learning alignment vectors among geometric sketches, which are then used to transform the full data sets to remove tissue-specific differences (**Methods**), decreases integration time of 534,253 human immune cells from hours to minutes while achieving comparable integration quality (**Supplementary Fig. 6**).

### Geometric Sketching Accelerates scRNA-seq Data Integration

In addition to being efficient by itself, geometric sketching can also accelerate other downstream algorithms for scRNA-seq analysis. One such problem involves integration of multiple scRNA-seq data sets across different batches or conditions (Butler et al., 2018; Haghverdi et al., 2018; Hie et al., 2018; Korsunsky et al., 2018). Here, we consider a novel approach to accelerating scRNA-seq data integration by applying the integration algorithm only to geometric sketches instead of the full data sets. Then, we use the integrated values of the sketch to learn a nonlinear transformation that is applied to the full data set to place it on the same integrated landscape (**Methods**). Since the integration step is more computationally intensive than the latter interpolation step, our geometric sketch-based integration offers a speedup that becomes especially dramatic when integrating large numbers of cells. Moreover, because geometric sketching better preserves rare transcriptional states, as demonstrated above, rare cell types are also more likely to be integrated during the procedure compared to using sketches from other sampling approaches.

We applied geometric sketch-based acceleration to two existing algorithms, Scanorama (Hie et al., 2018) and Harmony (Korsunsky et al., 2018), for scRNA-seq data integration (**Figure 8**). However, we note that our acceleration procedure is agnostic to the underlying integration method and can easily interface with similar algorithms (Butler et al., 2018; Haghverdi et al., 2018). We benchmarked the runtime improvement using geometric sketching on a data set of 534,253 human immune cells from two different tissues (umbilical cord blood and adult bone marrow). On this data, Scanorama and Harmony require 2.1 and 1.9 hours of computation, respectively, to obtain integrations that remove tissue-specific differences. In contrast, the integration procedure with geometric sketching (which includes finding the geometric sketches, integrating the sketches, and then transforming the full data sets based on the sketches) requires just 8 minutes of computation with either Scanorama or Harmony. Moreover, using geometric sketching-based acceleration has integration performance comparable to the full integration (**Figure 8**) and better than sketch-based integration using other sampling strategies (**Supplementary Fig. 6**), providing yet another example of how geometric sketching can be used to accelerate other algorithms for large-scale scRNA-seq analysis.

## DISCUSSION

Geometric sketching provides an efficient algorithm for obtaining subsamples of large scRNA-seq data sets such that the subsample contains as much of the transcriptional heterogeneity from the original data set as possible. Our algorithm’s sketches require less bandwidth to transfer and can be more easily shared among researchers. Geometric sketches can be inputted into computationally intensive downstream analysis tools designed for smaller data sets, including those that learn complex low-dimensional embeddings (Ding et al., 2018), 2-dimensional visualization coordinates (Cho et al., 2018; McInnes and Healy, 2018), or that fit complex models for a variety of tasks including pseudo-temporal trajectory analysis (Qiu et al., 2017), rare cell type discovery (Grün et al., 2015; Jiang et al., 2016), gene regulatory network reconstruction (Van Dijk et al., 2018), or robust differential expression analysis (Kharchenko et al., 2014). While our method does not distinguish between transcriptional structure due to biological or technical variation (e.g., batch effects), our sampling algorithm could be applied separately to data sets from different batches and then integrated or batch corrected using other methods (Butler et al., 2018; Hie et al., 2018).

Our work is distinct from but complementary to techniques that aim to find representative summaries of gene expression within clusters of cells (Baran et al., 2018; Iacono et al., 2018; Saunders et al., 2018), which output aggregate expression profiles that are not observed in the original data set. Applying geometric sketching as a preprocessing step would mostly likely accelerate these complex methods for gene expression aggregation while preserving representation of rare transcriptional states. Moreover, we note that because all of the elements within a geometric sketch correspond to actual observations from the original data, researchers have the flexibility to apply any existing downstream method designed for single-cell RNA-seq data sets, unlike methods that modify the gene expression values.

We note that our algorithm should be used in conjunction with other tools for scRNA-seq quality control. To limit artifacts arising due to dropout and data sparsity, it is common to apply a minimum unique gene cutoff, which we also do in our experiments; filtering steps with a linear time complexity in the size of the data set are unlikely to be a substantial bottleneck for single-cell methods. Another potential artifact common to droplet-based scRNA-seq experiments are doublets, which, due to their more complex transcriptional signatures, may also be more likely to appear within a geometric sketch. Many methods have been developed for computational doublet detection (DePasquale et al., 2018; Kang et al., 2018; McGinnis et al., 2018; Wolock et al., 2018), which can be applied to the sketch to remove these potential sources of confounding variation. We also note that more advanced quality control methods, including those for normalization (Bacher et al., 2017; Lun et al., 2016b; Vallejos et al., 2017), highly variable gene filtering (Yip et al., 2018), and imputation (Van Dijk et al., 2018; Li and Li, 2018; Ronen and Akalin, 2018) can naturally be applied to a geometric sketch before further analysis.

While it is possible for individuals to download large data sets and independently run geometric sketching, we envision laboratories that generate large-scale single-cell omics data sets would also compute and provide geometric sketches alongside the full data. These sketches would then be available to download for users with more limited computational resources or those wishing to run quick exploratory analyses on a subset of the data. In this spirit, we have computed small geometric sketches of a number of large, publicly-available scRNA-seq data sets containing hundreds of thousands of cells or even millions of cells, which are available for download at http://geosketch.csail.mit.edu. We also provide implementations of geometric sketching and the other sampling algorithms used in our benchmarking experiments in the geosketch Python package (https://github.com/brianhie/geosketch). Finally, we note that our techniques can be applied beyond single-cell transcriptomics, or even biological data sets, to any setting in which compact, geometric summaries of the data would prove useful.

## ACKNOWLEDGEMENTS

We thank Y.W. Yu for discussion and feedback. B. Hie and H. Cho are partially supported by NIH grant R01GM081871 (to B. Berger). A preliminary version of this manuscript has been accepted for presentation at the 2019 Annual International Conference on Research in Computational Molecular Biology (RECOMB).

## AUTHOR CONTRIBUTIONS

All authors conceived the algorithm. B. Hie and H. Cho performed the computational experiments. B. Bryson performed the scRNA-seq experiments. B. Berger led the research. All authors wrote the manuscript.

## METHODS

### Geometric Sketching Problem

We first give a mathematical formulation of the sketching problem to elucidate the theoretical insights underlying our approach. Let 𝒳 = {x_1_, …, x_*n*_} be a representation of a single-cell data set, consisting of *m*-dimensional measurements x_*i*_ ∈ ℝ^*m*^ from *n* individual cells. In the case of very large *n* (e.g., millions of cells) (Macosko et al., 2015; Zheng et al., 2017), it is often desirable to construct a *sketch* 𝒮 ⊂ 𝒳 (i.e., a downsampled data set), which can be more easily shared with other researchers and be used to quickly understand the salient characteristics of 𝒳 without paying the full computational price of analyzing 𝒳.

Drawing insight from computational geometry, we measure the quality of a sketch 𝒮 with respect to a data set 𝒳 via the *Hausdorff distance d*_*H*_ (Hausdorff, 1937) defined as

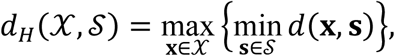

where *d* denotes the distance function of the underlying metric space (i.e., a notion of dissimilarity between two cells). Intuitively, *d*_*H*_ measures the distance of the cell in the original data set that is farthest away from any of the cells included in the sketch. The lower this distance, the more comprehensively our sketch covers the original data set.

We are interested in developing an efficient algorithm for obtaining 𝒮 of a predetermined size *k* (i.e., |*S*| = *k*) that minimizes *d*_*H*_(𝒳, 𝒮). A key property of our approach is that it is *agnostic to local density of data points*, since only the maximum distance is taken into account. As a result, our sketches more evenly cover the space of gene expression spanned by the original data set. In contrast, approaches based on uniform sampling or distance-based sampling [e.g., *k*-means++ (Arthur and Vassilvitskii, 2007)] are biased toward selecting more cells in densely populated regions at the expense of other regions of interest with fewer data points, as we demonstrate in our experiments.

### Theoretical Connection to Covering Problems

Our problem of finding a high-quality sketch 𝒮 of size *k* that minimizes *d*_*H*_(𝒳, 𝒮) is closely related to the concept of covering numbers in information theory and combinatorics. Informally, *internal covering number* is defined as the smallest number of equal-sized shapes (e.g., spheres or boxes) centered at individual data points that, together, “cover” all points in a data set. As we show in Lemma 1 in the **Supplementary Text**, the *minimum radius* for covering spheres that gives an internal covering number of at most *k* on a given data set is in fact equal to the optimal Hausdorff distance achievable by a sketch of size *k*. An important insight given by this observation is that the problem of finding a high-quality sketch reduces to finding a minimum-cardinality cover of a data set given a certain radius. In particular, if one were to have access to an oracle that could find the optimal covering of a data set for any radius, our problem could be solved by finding the minimum radius that gives the desired number of covering spheres (e.g., via binary search). Unfortunately, finding the minimum-cardinality cover is NP-complete (Attali et al., 2016), and although algorithms for a variety of simplified settings have been studied (Ahn et al., 2011; Alt et al., 2006; Chan and Hu, 2015; Chvatal, 1979), none scales to the high-dimensional and large-scale data that we need to handle in single-cell genomics. Given the hardness of the covering problem, we aimed to devise an approximate covering algorithm that readily scales to large-scale single-cell data while maintaining good sketch quality.

### Our Geometric Sketching Algorithm

At the core of our geometric sketching algorithm is a *plaid covering*, which approximates the geometry of the given single-cell data as a union of equal-sized boxes. To enable scalability to large data sets, we restricted our attention to covering the data points with a simple class of covering sets—plaids—whose structure is amenable to fast computation. Formally, we define a length-𝓁 *plaid cover* 𝒞 of a data set 𝒳 as a collection of points c_1_, …, c_*k*_ ∈ ℝ^*m*^ such that:

i. Either *c*_*ij*_ = *c*_*i*_′_*j*_ or |*c*_*ij*_ – *c*_*i*_′ _*j*_| ≥ 𝓁 for all *i, i*′ ∈ [*j*] and *j* ∈ [*m*], and
ii. 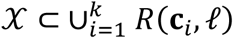, where *R*(c_*i*_, 𝓁) = [*c*_*i*1_, *c*_*i*1_ + 𝓁] ×… × [*c*_*im*_, *c*_*im*_ + 𝓁].

Intuitively, 𝒞 represents a collection of *m*-dimensional square boxes of side length 𝓁 covering 𝒳 that can be generated by placing a grid (with potentially uneven intervals) over the space and selecting a subset of grid cells. An example plaid cover is illustrated in **Figure 1**. Our greedy algorithm for finding a plaid cover of a given data set is shown in **Algorithm 1**. It can be shown that the plaid cover found by our algorithm uses the smallest number of intervals in each coordinate (see **Supplementary Text** for proof), although it may not achieve the smallest cardinality overall.

The main intuition behind our choice of plaid pattern is that it generalizes grid-based approximation of geometric shapes while maintaining computational efficiency in assigning points to their respective covering box. Note our plaid covering algorithm has time complexity in each dimension of *O*(*n* log *n*) in general—the main bottleneck being the sorting of each coordinate—and uses *O*(*n*) space. In practical scenarios where each coordinate requires only a small constant number of intervals to cover, we achieve *O*(*n*) time complexity by taking linear scans to find the next interval without sorting. This is a substantial improvement over other approaches for tackling the covering problem, which typically require *O*(*n*^2^) time for all pairwise distance calculations. A greedy approach to building a cover could require only *O*(*kn*) pairwise distance calculations where *k* is the number of covering objects (Chvatal, 1979), yet *k* is still typically much larger than log *n* for our applications in single-cell analysis.

#### Algorithm 1: Greedy Plaid Cover

~~~
**Data:** Data set 𝒳 = {x_1_, …, x_*n*_} where x_*i*_ ∈ ℝ^*m*^, length 𝓁
**Result:** Length-𝓁 plaid cover 𝒞 of 𝒳
y_*i*_ ← 0 ∈ ℝ^*m*^, ∀*i* ∈ [*n*]
**for** *j* ∈ [*m*] **do**
        z_1_, …, z_*n*_ ← **Sort**({*x*_1*j*_, …, *x*_*nj*_}) /* In ascending order. */
        *p* ← 1
        **while** z_*p*_ + 𝓁 < z_*n*_ **do**
               Find smallest *i* > *p* where z_*p*_ + 𝓁 < z_*i*_
               *y*_*i*_′_*j*_ ← z_*p*_, ∀*i*′ ∈ {*p*, …, *i* – 1}
               *p* ← *i*
        **end**
        *y*_*i*_′_*j*_← z_*p*_, ∀*i*′ ∈ {*p*, …, *n*}
**end**
**return** {y_1_, …, y_*n*_} /* Only unique points are returned. */
~~~

The cardinality of the cover returned by our plaid cover algorithm generally decreases as the length parameter 𝓁 increases, although pathological cases that deviate from this pattern exist. We empirically confirmed the near-monotonic relationship between number of covering boxes and 𝓁 on all our single-cell benchmark data sets (**Supplementary Fig. 7**). Based on this observation, we perform binary search (with graceful handling of potential exceptions) to find the value of 𝓁 that approximately produces a desired number of covering boxes. By default, we choose the same number of boxes as the desired sketch size *k*. A sketch is then constructed by sampling the boxes in a plaid cover and choosing a point at random from each box. Theorem 1 provided in **Supplementary Text** provides a theoretical insight into the quality of a sketch obtained via plaid covering. In particular, it gives a bound on the optimal Hausdorff distance relative to the solution obtained by plaid covering.

In order to reduce the dimensionality of the problem for scalability as well as robustness to noise, we first project the data down to a relatively low-dimensional space (100 dimensions for single-cell data) using a fast random projection-based singular value decomposition (SVD) (Halko et al., 2011) before applying our plaid covering algorithm. We note that much work has been done in obtaining algorithms for computing an approximate SVD of very large data sets with provable bounds on approximation error that are also highly efficient in runtime and memory (Halko et al., 2011; Ross et al., 2008); obtaining the top 100 principal components (PCs) of our largest benchmark data set with 665,858 cells requires about 10 minutes of additional computation time with linear-time scalability in the size of the data set (**Supplementary Fig. 8**).

### Geometric Sketching Algorithm Parameters

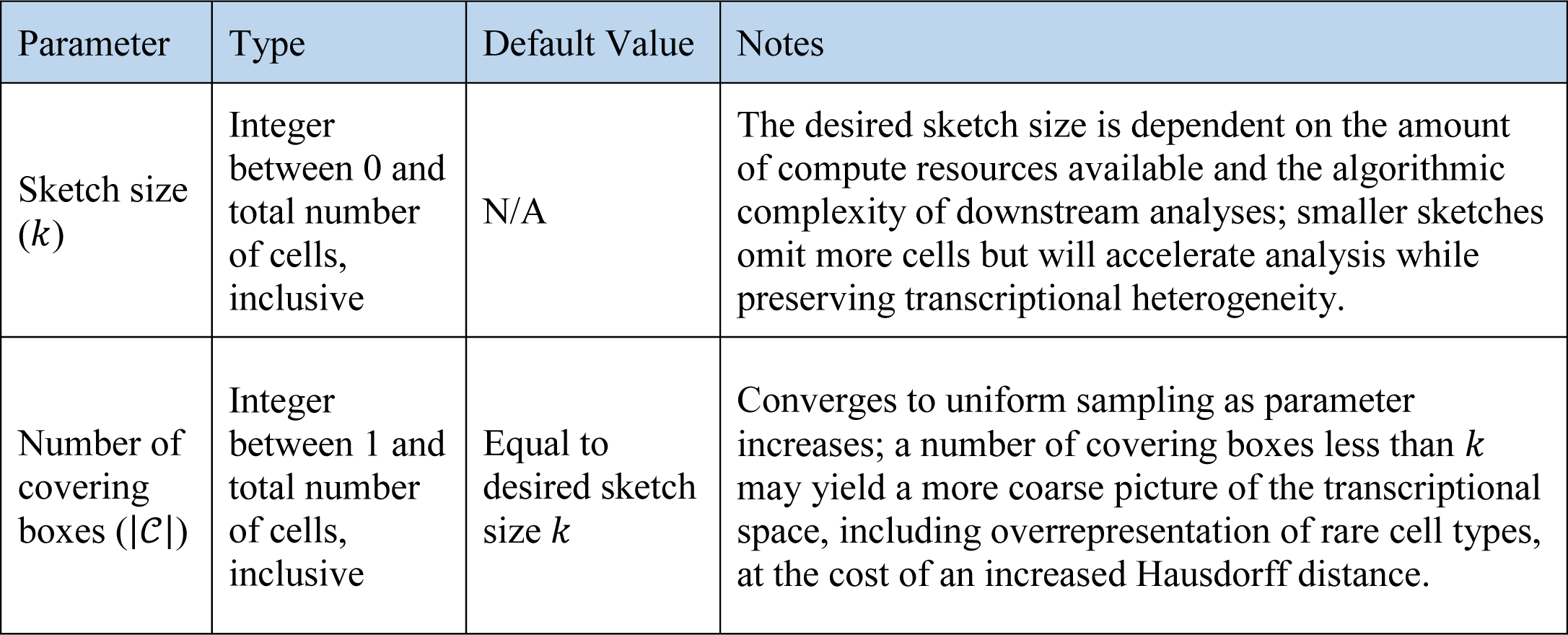

### Baseline Sampling Methods

We benchmark our algorithm against a number of existing sampling methods:

i. *Uniform sampling* returns a random sample of the cells, where every cell is given equal probability. We use the random choice function provided by the numpy Python package (Oliphant, 2006).
ii. *Spatial random sampling* (SRS) (Rahmani and Atia, 2017b) first projects the data points onto the unit hypersphere, then each sample is obtained by uniformly sampling a point on the unit hypersphere and selecting the closest point in the projected data set according to the cosine distance.
iii. *k-means++ sampling* (Arthur and Vassilvitskii, 2007) randomly chooses an initial sample, then repeatedly samples the next point by giving each point a weight proportional to the minimum distance from previous samples. This procedure continues until the desired number of samples have been obtained. We used the *k*-means++ implementation from the scikit-learn package (Pedregosa and Varoquaux, 2011).

We also run our experiments for SRS and *k*-means++ sampling using the same lower dimensional embeddings (top 100 PCs) used as input to geometric sketching.

Instead of first sampling the data and then clustering, it may be possible to do an expensive clustering step on the full data first, and then sample from those clusters. We note that, in addition to its lack of scalability, this procedure does not guarantee even coverage of the transcriptional space but is dependent on the assumptions of different clustering algorithms and is also sensitive to the choice of clustering parameters. We apply two common clustering strategies, *k*-means clustering (Steinhaus, 1956) and Louvain community detection (Blondel et al., 2008), for this purpose and compare to our method (**Supplementary Fig. 11**). Louvain clustering followed by sampling is the approach taken by the dropClust pipeline (Sinha et al., 2018).

### Benchmark Data Sets

We used the following data sets for our benchmarking experiments:

i. *293T and Jurkat mixture*. We obtained a mixture of 293T cells and Jurkat cells from 10X Genomics (Zheng et al., 2017) containing a much smaller number of 293T cells than Jurkat cells, where cell types are computationally inferred based on consensus clustering and marker genes. We removed cells below a cutoff of 500 unique genes, normalized each cell by the total expression and reduced the dimensionality to 100 PCs. The resulting data contained 4,185 cells in total.
ii. *Peripheral blood mono-nuclear cells (PBMCs)*. We obtained a data set of PBMCs from 10X Genomics (Zheng et al., 2017) and used the computationally curated cell type labels as well as the cell quality filtering steps from the original study. We then normalized each cell by the total expression and reduced the dimensionality to 100 PCs. The resulting data contained 68,579 cells.
iii. *Adult mouse brain*. We obtained scRNA-seq data from different regions of the mouse brain from Saunders *et al*. (Saunders et al., 2018) and used the computationally curated cell type labels as well as the cell quality filtering steps from the original study, including removal of doublet and outlier cells. We then normalized each cell by the total expression and reduced the dimensionality to 100 PCs. The resulting data contained 665,858 cells.
iv. *Developing and adolescent mouse central nervous system (CNS)*. We obtained scRNA-seq data from different regions of the mouse CNS from Zeisel *et al*. (Zeisel et al., 2018), removed cells below a cutoff of 500 unique genes, and used the computationally curated cell type labels and additional cell quality filtering steps from the original study. We then normalized each cell by the total expression and reduced the dimensionality to 100 PCs. The resulting data contained 465,281 cells.

### Robust Hausdorff Distance Computation

The classical Hausdorff distance (HD) (Hausdorff, 1937), according to our problem formulation, is computed as 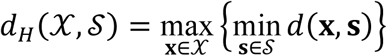 where 𝒳 denotes the point set corresponding to the full scRNA-seq data set and 𝒮 denotes the point set corresponding to a sketch, where 𝒮 ⊆ 𝒳. Because the classical HD measure is highly sensitive to even a few number of outliers (Huttenlocher et al., 1993; Sim et al., 1999), we use a robust HD measure proposed by Huttenlocher et al. called the partial HD measure, defined as 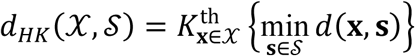 where 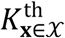 denotes the *K*^th^ largest value; partial HD requires a parameter *q* = 1 – *K*/|𝒳| between 0 and 1, inclusive, which is equivalent to classical HD when *q* = 0 (Huttenlocher et al., 1993). We set *q* = 1e-4, which obtains a measurement that is very close to the value obtained by classical HD but is robust to the most extreme outliers. We achieved similar results for different values of *q* (**Supplementary Fig. 9**).

### Data Visualization

To visualize the subsampled data based on different sampling methods, we used a 2-dimensional *t*-distributed stochastic neighbor embedding (*t*-SNE) with a perplexity of 500, a learning rate of 200, and 500 training iterations. We used the implementation provided by the Multicore-TSNE Python package (https://github.com/DmitryUlyanov/Multicore-TSNE).

### Simulation Analysis of Data with Known Volume and Density

To obtain data sets for which the volume (transcriptional diversity) and density of each cell type is known *a priori*, we considered different ways to duplicate and transform a data set composed entirely of 293T cells (Zheng et al., 2017). To obtain a data set with clusters of equal volume and variable density, we uniformly subsampled the 293T cells by a factor of 10 or 100 to create two new clusters, where each new cluster is translated such that none of the clusters overlap.

Likewise, to obtain a data set with clusters of equal number of data points but variable volume, we projected down to 3-dimensions using PCA and rescaled the components by a factor of 10 or 100 to create two new clusters, which are similarly translated to avoid overlap. Using a lower dimensionality in our simulations allowed us to better reason about the expected change in volume and is close to the effective fractal dimension of the data set (**Supplementary Fig. 10**). We then sketched these data sets and assessed the density-dependence by computing the Kullback-Leibler (KL) divergence 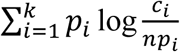, where *p*_*i*_ denotes the normalized volume of cell type *i* such that ∑_*i*_ *p*_*i*_ = 1, *c*_*i*_ denotes the number of cells in cluster *i, k* is the number of clusters, and *n* is the total number of cells in the data set. Lower values of the KL divergence indicate a sampling that better reflects the volume of each of the clusters.

### Differential Entropy of Cell Types

To obtain a rough estimate of the transcriptional variability represented by each cell type *i*, we fit a multivariate Gaussian distribution to each cell type to obtain an estimate of the covariance 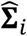; we then computed the differential entropy 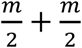 In 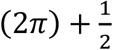 In 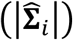 where *m* is the dimensionality of the data. We fit the distribution using the GaussianMixture class from scikit-learn (Pedregosa and Varoquaux, 2011).

### Clustering Analysis

We quantify the ability for clustering analyses on a subsample of a full data set to recapitulate a set of “ground truth” labels, in this case, the cell type labels assigned by the original study authors.

For the Louvain clustering analysis, we constructed the nearest neighbors graph on which we applied the Louvain community detection algorithm (Blondel et al., 2008). We use the graph construction and Louvain implementation with default parameters provided by scanpy (Wolf et al., 2018), which leverages the louvain-igraph package (https://github.com/vtraag/louvain-igraph). Louvain cluster labels were applied to the full data set based on the most common label of the five nearest neighbors within the sketch (ties broken randomly). We quantified agreement between the unsupervised cluster labels and the previous study labels using the adjusted mutual information (AMI) score (Vinh et al., 2010) implemented by the scikit-learn Python package (Pedregosa and Varoquaux, 2011) based on a resampled data set where the relative frequencies of the ground truth clusters are set to uniform to equally consider the clusters regardless of their abundance in the full data set. We refer to this metric as balanced AMI (BAMI). The correction factor in AMI for chance agreement is updated accordingly to account for the balanced distribution. We repeat the analysis using three different Louvain resolution parameters (0.5, 1, and 2) and take the maximum BAMI score across these parameter settings for each sampling algorithm.

### Immune Cell Analysis

254,941 cells from umbilical cord blood were obtained from the Human Cell Atlas (https://preview.data.humancellatlas.org). The data set was filtered for cells containing more than 500 unique genes, normalized by the total expression for each cell, and then natural log transformed after adding a pseudo-count of 1. Data was projected to 100 PCs using the randomized PCA implementation provided by the fbpca Python package (https://github.com/facebook/fbpca). Unsupervised clustering was performed by running the Louvain community detection algorithm with the default parameters (resolution of 1, 15-nearest neighbors graph) of the scanpy framework (Wolf et al., 2018). Marker genes for inflammation were selected using a nominal AUROC cutoff of 0.9 for separation of the inflammatory cluster from the remaining clusters of macrophages. Validation of marker genes in GM-CSF-versus M-CSF-polarized macrophages using a one-sided Welch’s *t-*test (for unequal population sizes) using the scipy Python package (Oliphant, 2007).

### Macrophage Polarization scRNA-seq Experiment and Analysis

scRNA-seq data of M-CSF-derived macrophages was obtained from the study by Hie *et al*. (2018). We repeated the same experiment but instead polarized with an inflammatory stimulus, GM-CSF. Human monocytes were isolated from human buffy coats purchased from the Massachusetts General Hospital blood bank using a standard Ficoll gradient and subsequent CD14+ cell positive selection (Stemcell Technologies). Selected monocytes were cultured in ultra low-adherence flasks (Corning) for 6 days with RPMI media (Invitrogen) supplemented with 10% FBS (Invitrogen) and 50 ng/mL human GM-CSF (Biolegend). SeqWell analysis was performed as previously described (Gierahn et al., 2017). Briefly, after 6 days, cells were detached using trypsin, spun down, and counted. Approximately 12,000 cells were loaded on each array for each timepoint and condition to minimize doublet-loading. The arrays were sealed with a semi-permeable membrane prior to cell lysis and hybridization to single-cell beads. Beads were subsequently pooled for reverse transcription and whole transcriptome amplification.

Read alignment and transcript quantification were performed as in Macosko *et al*. (2015b). Briefly, raw sequencing data was converted to demultiplexed FASTQ files using bcl2fastq2 based on Nextera N700 indices corresponding to individual samples/arrays. Reads were then aligned to the hg19 genome using the Galaxy portal maintained by the Broad Institute for Drop-Seq alignment using standard settings. Individual reads were tagged according to the 12-bp barcode sequence and the 8-bp UMI contained in Read 1 of each fragment. Following alignment, reads were binned onto 12-bp cell barcodes and collapsed by their 8-bp UMI. Digital gene expression matrices for each sample were obtained from quality filtered and mapped reads, with an automatically determined threshold for cell count. Analysis was done on cells that were filtered for a minimum cutoff of 500 unique genes, normalized by the total expression of each cell, and then natural log transformed after adding a pseudo-count of 1.

### Runtime Benchmarking

We benchmarked the runtime of geometric sketching to obtain 20,000 cells from a data set of 1, 2.5, 5, or 10 million cells, where we obtained each full data set by resampling the cells from the mouse CNS data set (Zeisel et al., 2018) to reach the desired cell count. We timed the algorithm using Python’s time module. All experiments were done on a 2.30 GHz Intel Xeon E5-2650v3 CPU.

### Geometric Sketching-Accelerated Integration

We assume an integration function that takes in a list of data sets and returns modifications to the data sets that removes differences to due batch effect etc. Let X ∈ ℝ^*n*×*m*^ denote one of the data sets, X_𝒮_ ∈ ℝ^|𝒮|×*m*^ denote the subset of X obtained by geometric sketching, and 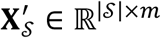 denote the modified version of X_𝒮_ returned by the integration function. Our goal is to apply a transformation to X that puts it into the same integrated space as 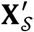. At a high level, we use a nearest-neighbors-based method to compute alignment vectors from X to X_𝒮_, we use Gaussian smoothing to combine these alignment vectors into translation vectors, and then we apply the translation to X to obtain an “integrated” full data set X^′^.

Formally, for each cell in X_𝒮_, we find its *k* nearest neighbors in X and we denote the set of all matches between a cell in X_𝒮_ and X as ℳ where |ℳ| = *k*|X_𝒮_|. Now we define the alignment vectors as the rows of the matrix 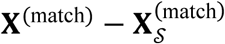 where the rows of 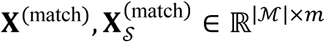 correspond to the pairs of matching cells in ℳ. We want to combine these alignment vectors to obtain our translation vectors, which we do using Gaussian smoothing. We compute weights via a Gaussian kernel as

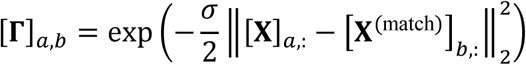

where Γ ∈ ℝ^*n*×|ℳ|^ and [·]_*a,b*_ denotes the value in the *a*^th^ row and *b*^th^ column of a matrix and [·]_*a*,:_ denotes the *a*^th^ row of a matrix. Finally, we construct the translation vectors as an average of the alignment vectors with Gaussian-smoothed weights, where

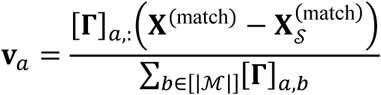

and we translate

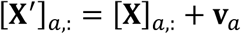

for all *a* ∈ [*n*] where [*n*] denotes the set of all natural numbers up to *n*. We repeat this for all data sets integrated by the “black-box” integration function; in our study, we used the Scanorama (Hie et al., 2018) and Harmony (Korsunsky et al., 2018) algorithms for integration.

We use geometric sketches of size 4000 (around 1% of the total data) and parameters *k* = 3 and *σ* = 15. We used Harmony version 0.0.0.9000 and Scanorama version 1.0. For all methods, we measured the runtime required for integration and translation, not including the initial PCA step for computing low dimensional embeddings (100 PCs). We quantify data set mixing by clustering the integrated embeddings using *k*-means, varying the number of clusters, and computing the average negative Shannon entropy normalized to a maximum value of 1 on the data set labels averaged across all clusters, an approach taken by recent work (Park et al., 2018).

### Data and software availability

We used the following publicly available data sets:

- 293T cells (Zheng et al., 2017) (https://support.10xgenomics.com/single-cell-gene-expression/datasets)
- 293T and Jurkat cell mixture (Zheng et al., 2017) (https://support.10xgenomics.com/single-cell-gene-expression/datasets)
- Human PBMCs (Zheng et al., 2017) (https://support.10xgenomics.com/single-cell-gene-expression/datasets)
- Developing and adolescent mouse CNS (Zeisel et al., 2018) (http://mousebrain.org)
- Adult mouse brain (Saunders et al., 2018) (http://dropviz.org)

Our code and data (including the above data sets) are available at http://geosketch.csail.mit.edu.

