## Supplementary Information for "Geometric Sketching Compactly Summarizes the Single-Cell Transcriptomic Landscape"

**Supplementary Materials for "Geometric Sketching Compactly Summarizes the Single-Cell Transcriptomic Landscape" by Hie, Cho, DeMeo, Bryson, and Berger**

Lemma 1

Let $\mathcal{X} = \{\mathbf{x}_1, \dots, \mathbf{x}_n\}$ be a representation of a single-cell data set, consisting of $m$-dimensional measurements $\mathbf{x}_i \in \mathbb{R}^m$ from $n$ individual cells. Let $d_H^*$ be the minimal Hausdorff distance $d_H(\mathcal{X}, \mathcal{S})$ obtained by a sketch $\mathcal{S} \subset \mathcal{X}$ where $|\mathcal{S}| = k$. Then, $d_H^* = N_{\text{int}}^{-1}(k)$, where $N_{\text{int}}^{-1}(k) \coloneqq \min\{r : N_{\text{int}}(\mathcal{X}, r) \leq k\}$.

Proof of Lemma 1

Since $d_H$ bounds the maximum distance of a data point from $\mathcal{S}$, placing a sphere of radius $d_H$ at every point in $\mathcal{S}$ gives a covering of $\mathcal{X}$, which implies $N_{\text{int}}(\mathcal{X}, d_H^*) \leq k$. Thus, $N_{\text{int}}^{-1}(k) \leq d_H^*$. If $N_{\text{int}}^{-1}(k) < d_H^*$, then there exists a cover with $k$ spheres of radius $d' < d_H^*$. Taking the center points of this cover as our sketch $\mathcal{S}'$, we obtain $d_H(\mathcal{X}, \mathcal{S}') \leq d_H^*$, a contradiction. Hence, $d_H^* = N_{\text{int}}^{-1}(k)$.

Theorem 1

Given a data set $\mathcal{X}$ of $n$ points in $m$ dimensions, let $N_{\text{plaid}}(\ell)$ be the number of boxes in the plaid cover returned by our algorithm as a function of length parameter $\ell$. Let $N_{\text{plaid}}^{-1}(k) = \inf\{\ell : N_{\text{plaid}}(\ell) \leq k\}$. Let $k$ be a desired sketch size and assume $k = N_{\text{plaid}}\left(N_{\text{plaid}}^{-1}(k)\right)$ for simplicity (if not take a nearby $k$ where this holds). Let $\mathcal{S}_{\text{plaid}}(k)$ be a sketch of size $k$ obtained by randomly choosing appoint from each box in the plaid cover. Let $d_H^*(k) = \min_{\mathcal{S}:|\mathcal{S}|=k} d_H(\mathcal{X}, \mathcal{S})$. Then, the following holds:

$$\frac{1}{2} N_{\text{plaid}}^{-1}(2^m \cdot k) \leq d_H^*(k) \leq d_H\left(\mathcal{X}, \mathcal{S}_{\text{plaid}}(k)\right).$$

Proof of Theorem 1



**Preprint. Work in progress.**

For the first inequality, Let $\mathcal{P} = P_1, P_2, \ldots, P_N$ be any covering by plaid sets of side length $2d_H^*(k)$, such that all covering sets contain at least one point. We show that $\mathcal{P}$ has cardinality at most $2^m k$.

Let $\mathcal{B}$ be a covering of $\mathcal{X}$ by $k$ balls $B_1, B_2, \ldots, B_k$, each with radius $d_H^*(k)$. The definition of $d_H^*$ ensures that such a covering exists. Define

$$I_{\mathcal{P}}(B_i) = \left|\left\{P_j : P_j \cap B_i \neq \emptyset\right\}\right|.$$

That is, $I_{\mathcal{P}}(B_i)$ is the number of sets in $\mathcal{P}$ that intersect $B_i$.

Because $\mathcal{P}$ and $\mathcal{B}$ are both covering sets, each plaid square in $\mathcal{P}$ is intersected by at least one ball in $\mathcal{B}$. Therefore,

$$|\mathcal{P}| \leq \sum_{i=1}^{k} I_{\mathcal{P}}(B_i).$$

On the other hand, we see that $I_{\mathcal{P}}(B_i)$ is bounded above by $2^m$, because any ball overlaps at most two plaid intervals in each dimension. Thus,

$$|\mathcal{P}| \leq 2^m k$$

as desired.

The second inequality is immediate, because $d_H^*(k)$ is an infimum of Hausdorff distances of all sets of size $k$ with $\mathcal{X}$, and $\mathcal{S}_{\text{plaid}}(k)$ is such a set.

Proof that Algorithm 1 is optimal in each dimension separately

Fix a dimension $d \in [n]$, and consider covering the projection $\pi_d(\mathcal{X}) = \{x_{1d}, x_{2d}, \ldots, x_{nd}\} \subset \mathbb{R}$ with a one-dimensional plaid cover of length $\ell$. Let $Q = \{q_1, \ldots, q_k\}$ be any such cover, and let $Y = \{y_1, \ldots, y_m\}$ denote the cover produced by our algorithm on iteration $d$. We show that $k \geq m$, i.e., $Y$ has the smallest size of any length-$\ell$ cover.



**Preprint. Work in progress.**

Assume without loss of generality that $q_1 < q_2 < \ldots < q_k$ and $y_1 < y_2 < \ldots < y_m$. Let $z_i$ denote the $i^{\text{th}}$-smallest element of $\pi_d(\mathcal{X})$. Our algorithm sets $y_1 = z_1$. We must have $q_1 \leq z_1$, or else $z_1$ is not covered by $Q$. Thus, $q_1 \leq y_1$. Proceeding inductively, we see that:

$$q_{i+1} \leq \min\{z_i : z_i > q_i + \ell\} \leq \min\{z_i : z_i > y_i + \ell\} = y_{i+1}$$

where the final equality holds because our algorithm defines $y_{i+1}$ exactly this way. Thus, we have $q_i \leq y_i$ for all $i \in 1, 2, \ldots, \min(k, m)$. If $|Q| \leq |Y|$, then $y_{m-1}$ and $y_m$ are both greater than all elements in $Q$. But because $Q$ covers all the points $z_i$, this implies that $y_m$ covers no points, a contradiction because our algorithm does not construct empty covering sets. Thus, we must have $|Q| \geq |Y|$, and because $Q$ is arbitrary, $Y$ has the smallest possible size.

Benchmarking against clustering-based analyses

In addition to comparing geometric sketching against other sampling methods, we also compared to a different approach (not typically used) in which a clustering step is first performed on the full data, which may be very expensive if the size of the data set is very large, and then a random subsampling is carried out by randomly selecting elements from each cluster. We test two clustering algorithms commonly applied to single cell data sets:

(i) *k-means clustering* [1] first clusters the data using a specified number of clusters, which we set to the square root of the data set size. Then, to obtain each sample, a cluster is chosen uniformly at random and a random point from the cluster is sampled uniformly. We initialize the *k*-means algorithm using *k*-means++ sampling, but note that using *k*-means++ itself as a sampling method is different from obtaining samples from the resulting clusters after iterative optimization.

(ii) *Louvain clustering* uses the Louvain community detection algorithm [2] to cluster the data; then, similar to the *k*-means-based sampling, a cluster is chosen uniformly at



**Preprint. Work in progress.**

random and then a cluster member is chosen also uniformly at random. We use Louvain resolution parameters of 1 or 3 [3], where higher resolutions tend to increase the number of clusters. This is the same algorithm used for "structure preserving sampling" in the dropClust pipeline [4].

Even in this more complex setting, our algorithm performs better at covering the full data set and is much more efficient than the other approaches; on the adult mouse brain data set [5], our algorithm samples 2% of the full data set in 3 minutes compared to around on hour on a single core for both Louvain clustering and *k*-means clustering (**Supplementary Fig. 11**).



**Preprint. Work in progress.**

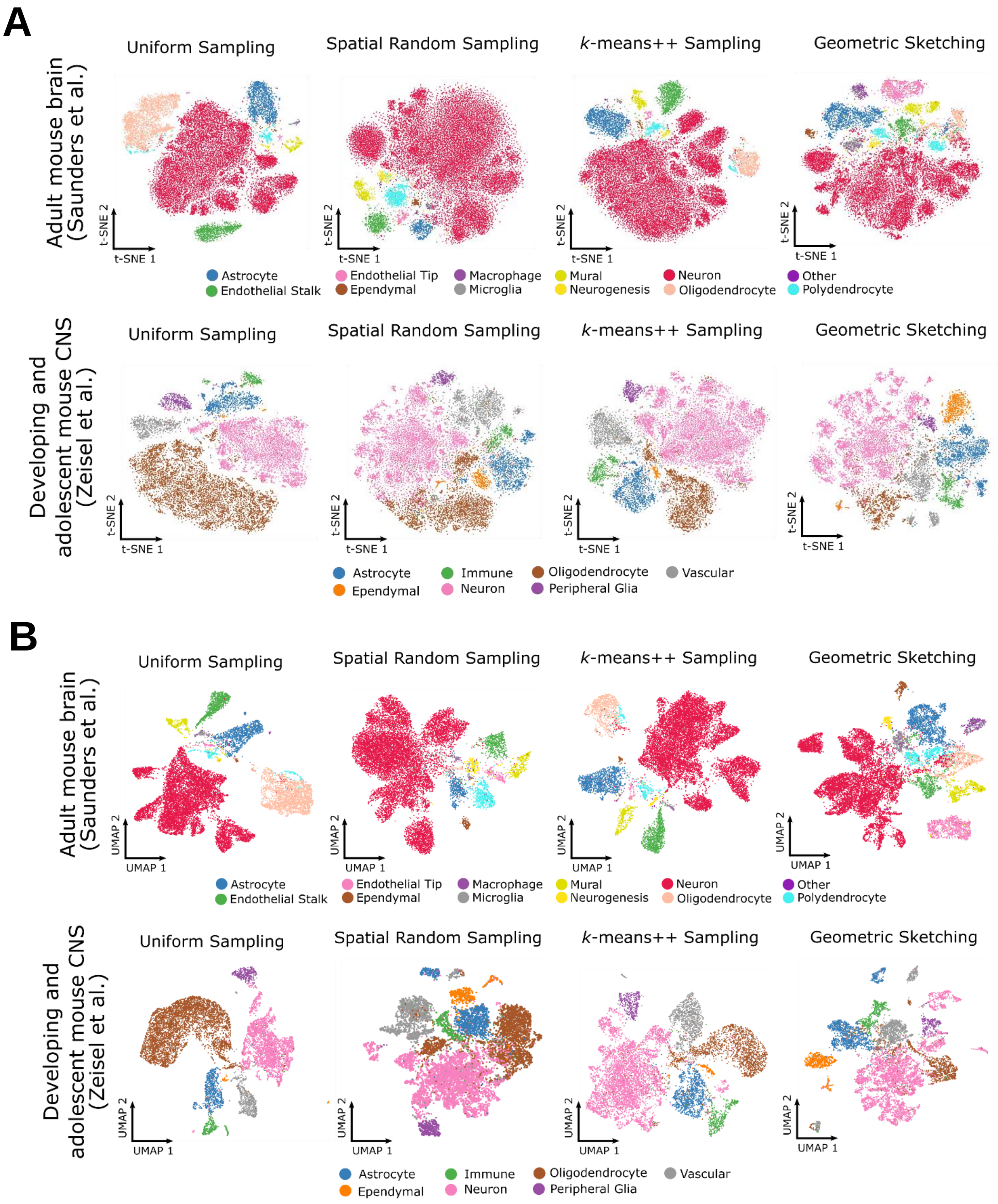


**Supplementary Figure 1: Visualizations of Different Sketches of Large-Scale scRNA-seq Data Sets**

Visualizations using (**A**) *t*-SNE and (**B**) UMAP of sketches containing 2% of the cells from the adult mouse brain [5] and from the developing and adolescent mouse CNS [6] using uniform random sampling, SRS, *k*-means++ and geometric sketching. Numbers of cells from each cell type are given in **Supplementary Tables 5-6**. Note that all data-dependent sampling methods underrepresent oligodendrocytes compared to uniform sampling, which is expected given the low transcriptional heterogeneity among oligodendrocytes as quantified by differential entropy (**Supplementary Tables 3-4**). While some sketches obtained by *k*-means++ sampling and SRS may appear similar to geometric sketches, they quantifiably preserver fewer rare cell types and have lower sketch quality as measured by the Hausdorff distance, which we show in our experiments.

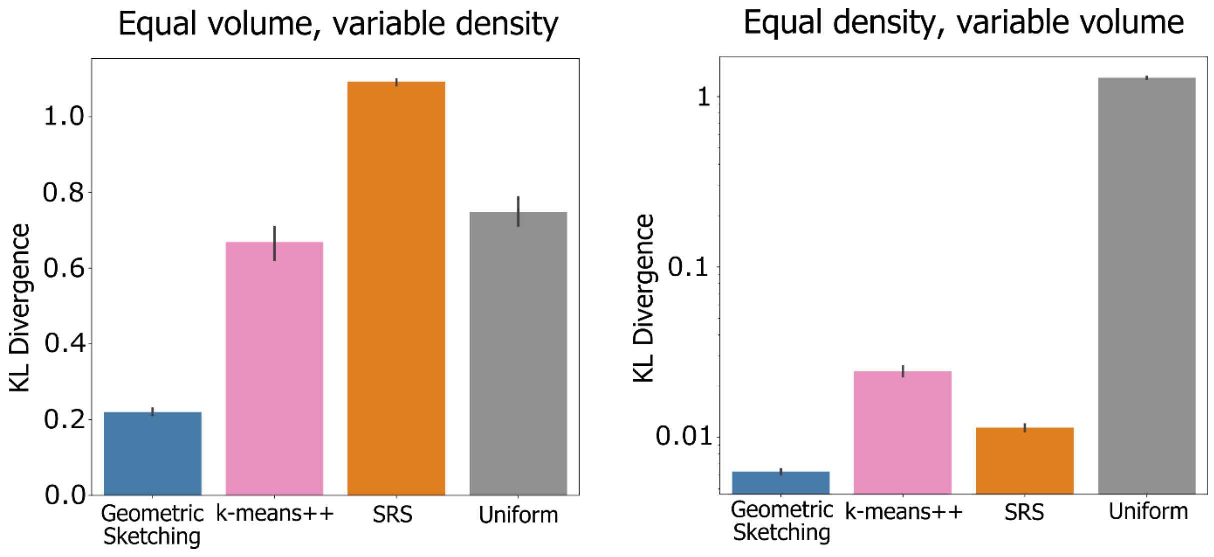


**Supplementary Figure 2: Sampling with Geometric Sketching Better Reflects Differences in Cluster Volume instead of Density**

Geometric sketching samples from clusters according to the volume of space occupied by each cluster. Bar height indicates means and error bars indicate standard error across 10 random seeds. The $y$-axis indicates the KL divergence of expected cluster representation based on known cluster volumes compared to observed cluster representation in the subsampled data; KL divergences for the equal density, variable volume experiment are plotted on a log scale. Closer to 0 is better (indicates less bias introduced by density). The data sets consist of clusters of equal volume but varying densities or clusters with equal numbers of cells but varying volumes.

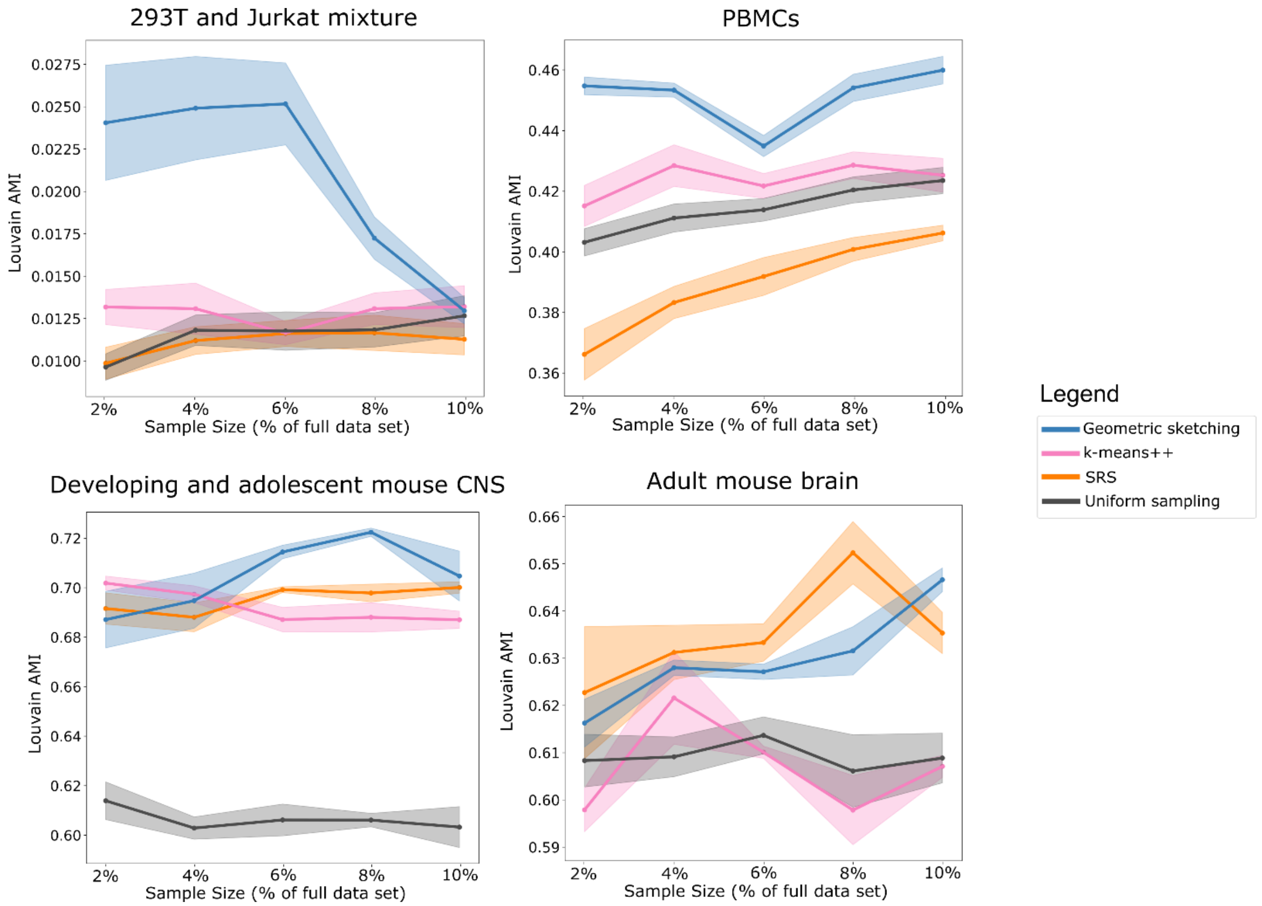


**Supplementary Figure 3: Rarest Cell Types Are More Represented within a Geometric Sketch**

We assessed overrepresentation of cell types within a sketch by computing the ratio of the observed number of cells over the expected number of cells (assuming uniform sampling probability) for each cell type; we then took the geometric mean of the ratios for the rarest half of all cell types within each data set. Geometric sketching consistently overrepresents rare cell types and does so more than other sampling strategies in almost all cases. Because we set the number of covering boxes equal to the desired sketch size, as the sketch size increases, the overrepresentation ratio with respect to uniform sampling will converge to unity.

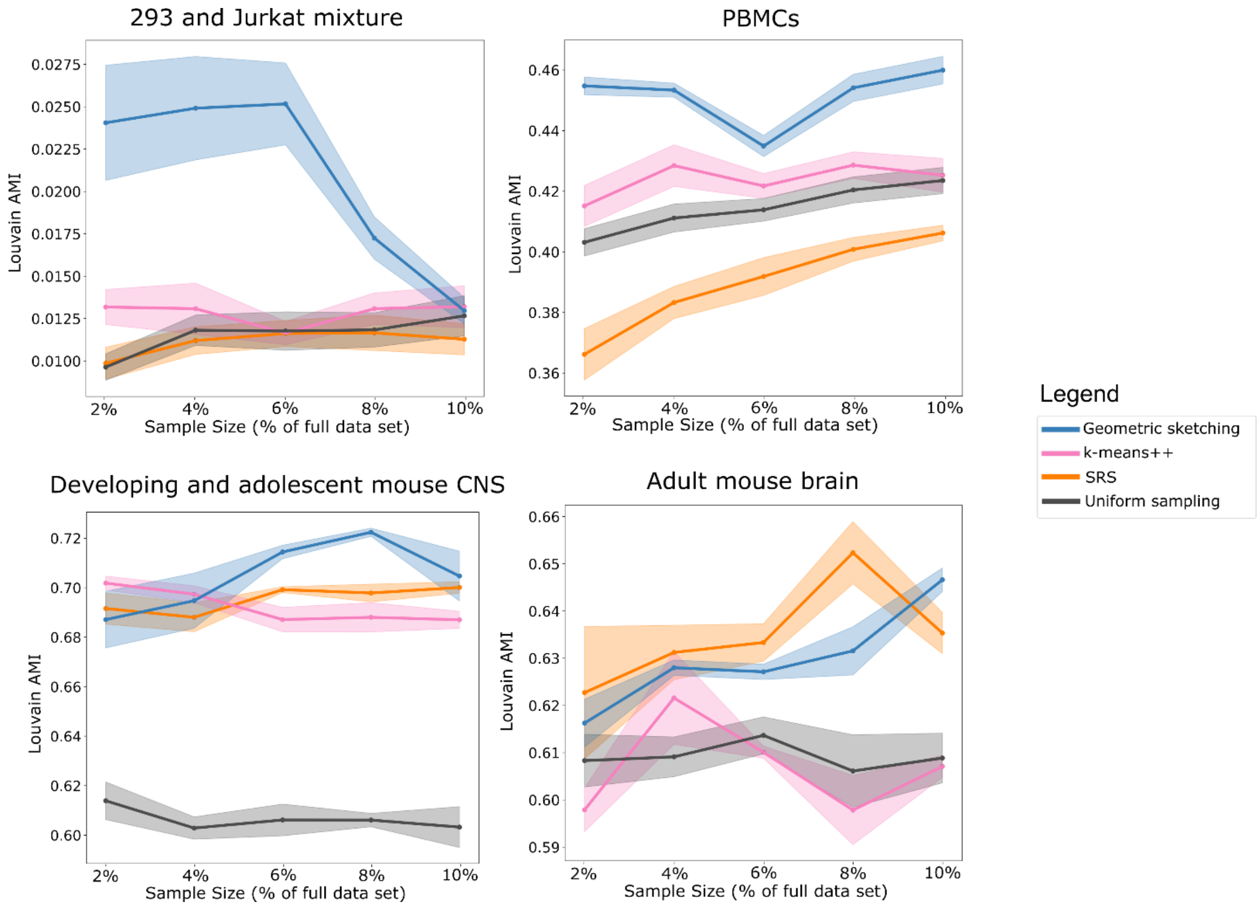


**Supplementary Figure 4: Unbalanced Measurement of Clustering Recapitulation of Biological Cell Types**

The same result as in **Figure 5** but without equal weighting of biological cell types. Louvain clustering was applied to a sketch, transferred to the full data set, and then measured for agreement with biological cluster labels using adjusted mutual information [7]. Unsupervised clustering of geometric sketches more consistently recapitulates biological cell types than clustering results obtained by uniform sampling and is comparable to or better than clusters of sketches from *k*-means++ and SRS.

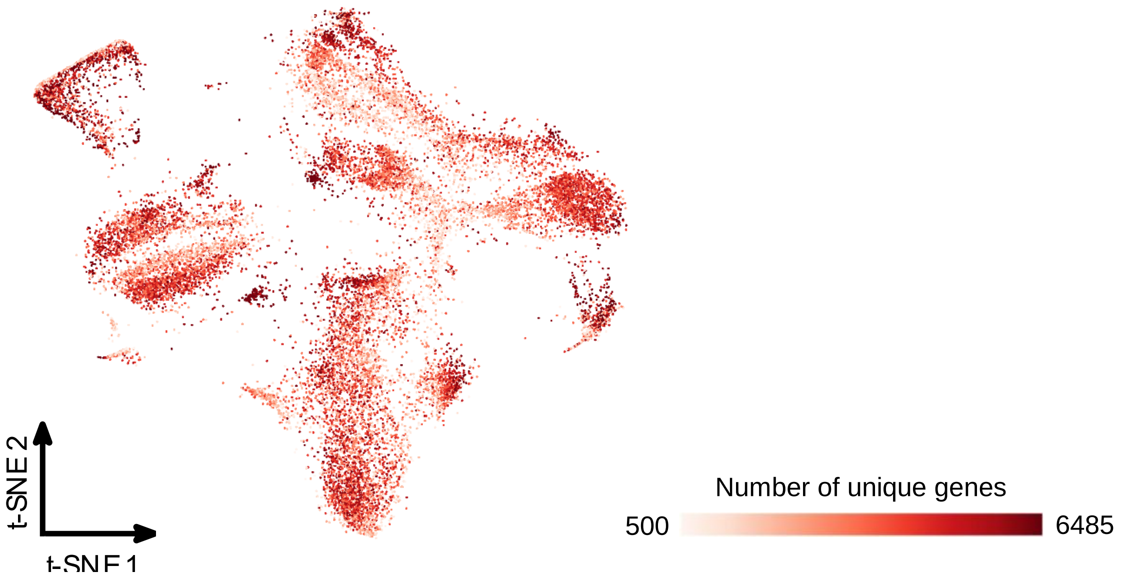


**Supplementary Figure 5: Unique Gene Heatmap of Umbilical Cord Cells**

Heatmap of the *t*-SNE embedded geometric sketch visualizing cells from human umbilical cord blood colored by the number of unique genes. Lighter red indicates higher levels of sparsity and darker red indicates lower levels of sparsity. The lowest number of unique genes in the data set was 500 and the highest was 6485 out of a total of 33,694 genes considered in the study.



**Preprint. Work in progress.**

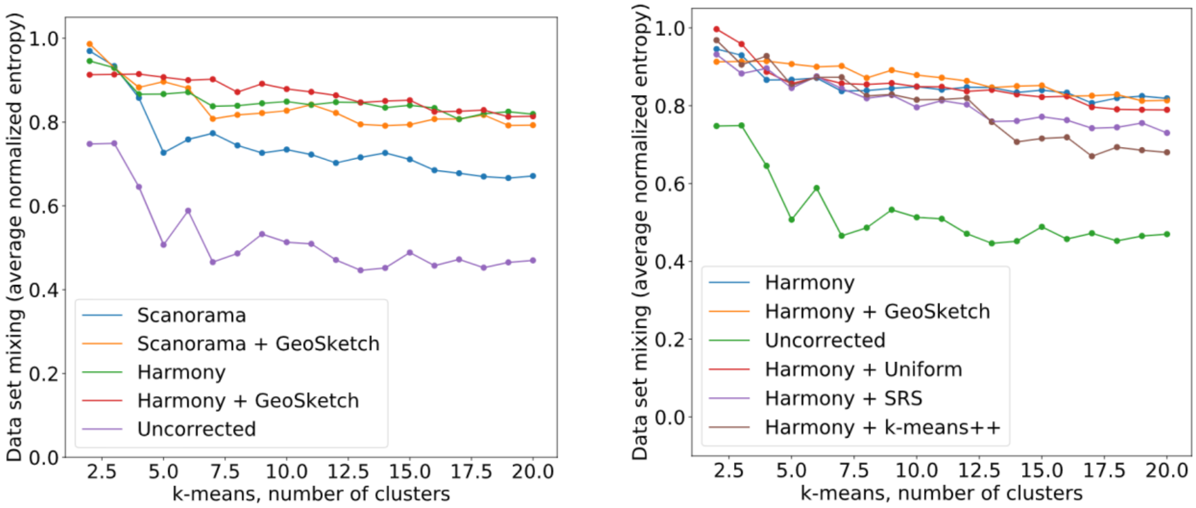


**Supplementary Figure 6: Integration Quality of Methods with and without Geometric Sketching-Based Acceleration**

Closer to 1 indicates more data set mixing within clusters; see **Methods** for description of our integration quality metric. Geometric sketching-based acceleration of integration methods yields integrations with comparable or better quality than applying the integration methods to the full data set. Both geometric sketching and uniform sampling have comparable integration quality, but based on our other results, it is likely that geometric sketching would better align rare cell types in addition to common cell types. Using SRS and *k*-means++ sampling produces worse integration quality.

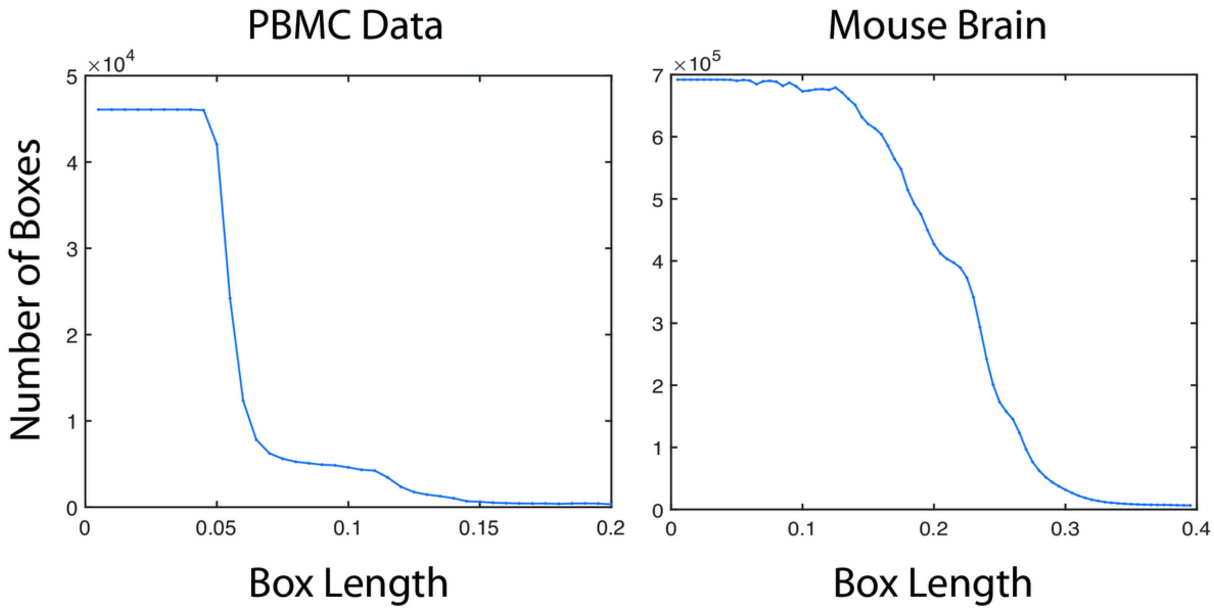


**Supplementary Figure 7: Near Monotonicity of Covering Boxes with Box Length**

Cardinality of plaid covering near-monotonically decreases with respect to the length parameter. For PBMC and adult mouse brain data sets, we plotted the number of boxes returned by our plaid covering algorithm as a function of box length provided as input. The overall monotonic relationship allows us to use binary search to find the length at which the plaid cover contains roughly the desired number of boxes.

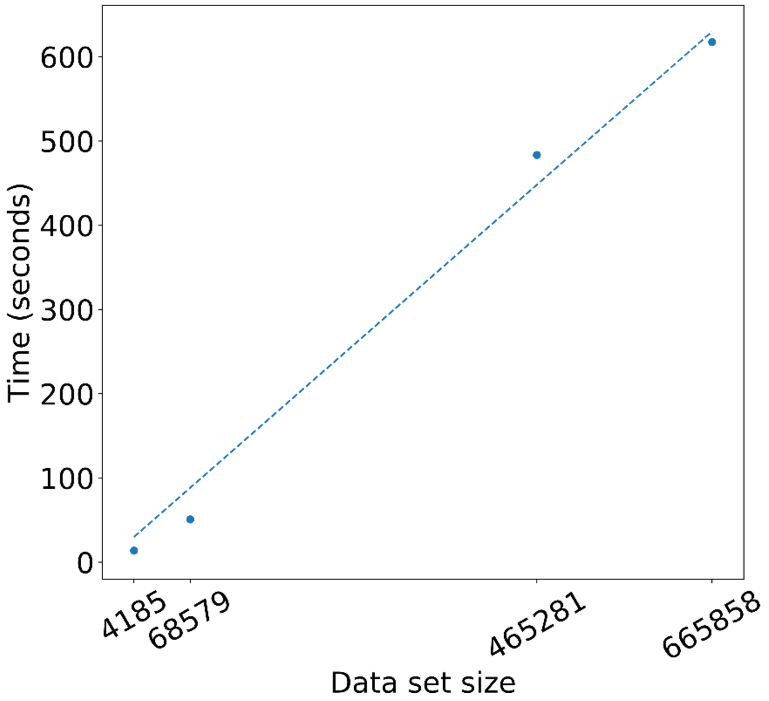


**Supplementary Figure 8: SVD Runtime versus Data Set Size**

The time required to learn a 100-dimensional representation of a scRNA-seq data set using a randomized SVD [8] scales linearly with the size of the data set and has reasonable scalability to large-scale scRNA-seq experiments in the future. Each point given in the above plot corresponds to the time taken to compute a 100-dimensional embedding on each of the four main benchmark data sets used in the study.

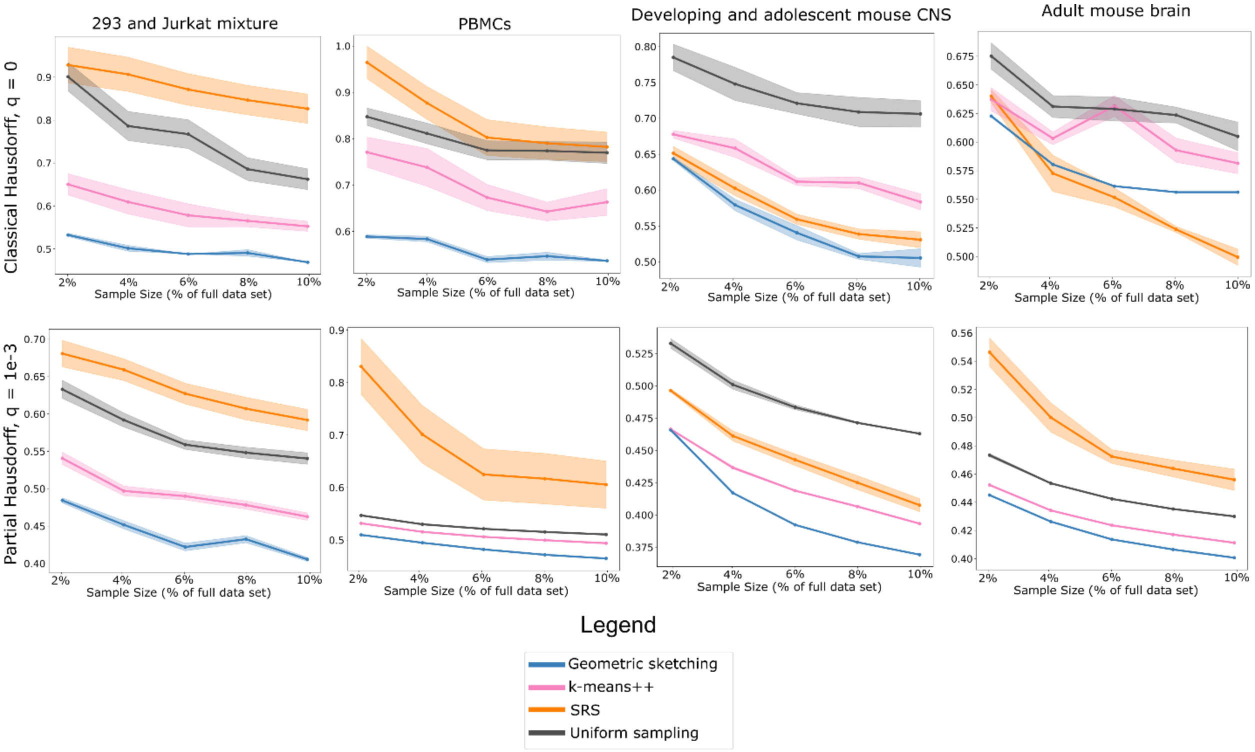


**Supplementary Figure 9: Partial Hausdorff Distance at Different Parameter Cutoffs**

We measured the partial Hausdorff distance at different values of the parameter $q$ (**Methods**), including $q$ = 1e-4 (**Figure 2**), $q$ = 1e-3 and $q = 0$ (the last corresponding to the classical Hausdorff distance). Geometric sketching outperforms all other sampling methods when measured with a robust, partial Hausdorff distance with positive $q$. Under the classical Hausdorff distance, geometric sketching also outperforms all other sampling methods in almost all cases except for larger sketches in the adult mouse brain data set due to a single outlier cell, but the anomalous cell was removed when computing more robust Hausdorff distance measures.



**Preprint. Work in progress.**

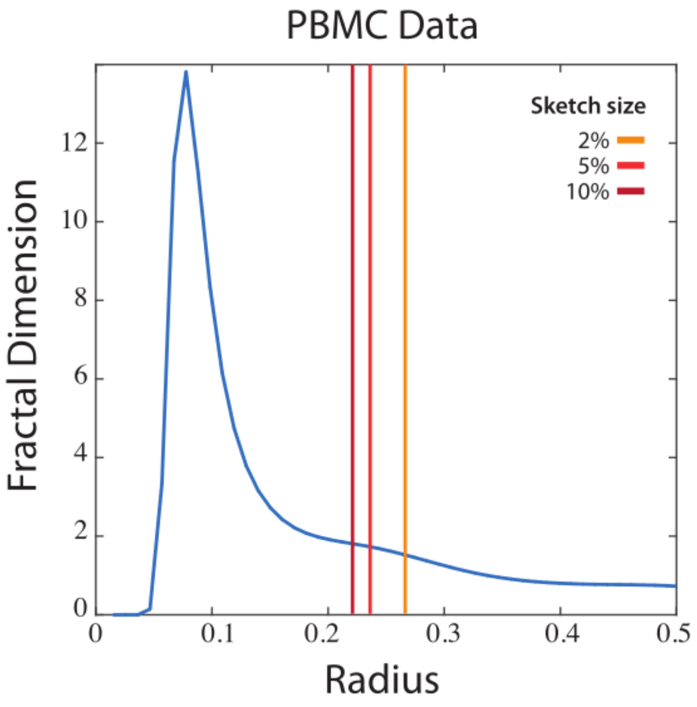


**Supplementary Figure 10: Low Fractal Dimension of Single-Cell Data**

On the PBMC data set, we computed the fractal dimension (averaged over the data points) at varying box lengths using the Chebyshev metric, which induces covering spheres that appear as boxes. Letting $N_r(x)$ be the number of data points covered by a sphere of radius $r$ centered at $x$, we define fractal dimension as $\log\left(N_{r_1}(x)/N_{r_2}(x)\right)/\log(r_1/r_2)$. Plot shows fractal dimension computed over 50 evenly spaced intervals between 0 and 0.5. Various vertical lines denote the radiuses that corresponds to the box size chosen by our geometric sketching algorithm when obtaining sketches containing different percentages of the overall data set. At the scale at which our geometric sketching operates, PBMC data displays a low fractal dimension of around 2.

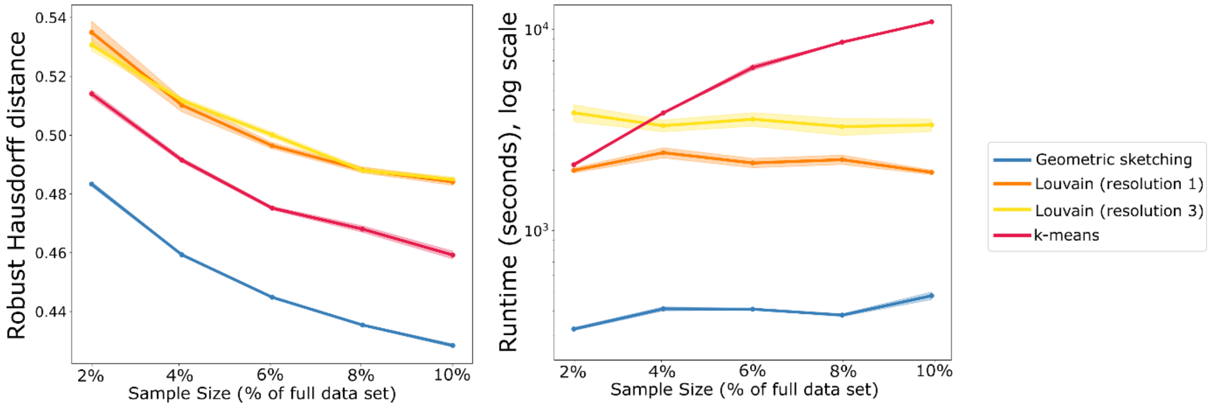


**Supplementary Figure 11: Comparison with Clustering-Based Sampling Methods**

Geometric sketching has comparable or improved performance over the much more computationally intensive strategy of clustering the full data set and then sampling within each cluster. On a data set of 665,858 cells from the developing and adolescent mouse CNS [6], geometric sketching has more even coverage of the data and is substantially faster than non-uniform sampling methods. The *y*-axis for the runtime plot is given on a logarithmic scale.



**Preprint. Work in progress.**

| Cell Type | Number of cells | % of total | Differential Entropy |
|---|---|---|---|
| 293T | 28 | 0.669056 | -461.66 |
| Jurkat | 4157 | 99.33094 | -270.88 |

**Supplementary Table 1**

Statistics for 293/Jurkat mixture data; for the differential entropy calculation, see **Methods**.

| Cell Type | Number of cells | % of total | Differential Entropy |
|---|---|---|---|
| CD14+ Monocyte | 3817 | 5.565844 | -228.419 |
| CD19+ B | 3306 | 4.820718 | -213.47 |
| CD4+/CD25 T | 2812 | 4.100381 | -238.942 |
| CD4+/CD45RA+/CD25- Naive T | 3126 | 4.558247 | -230.899 |
| CD4+/CD45RO+ Memory | 5859 | 8.543432 | -223.313 |
| CD4+ Helper T | 11445 | 16.68878 | -222.592 |
| CD56+ NK | 14112 | 20.57773 | -232.116 |
| CD8+/CD45RA+ Naive Cytotoxic | 21975 | 32.04334 | -232.351 |
| CD8+ Cytotoxic T | 1865 | 2.719491 | -219.693 |
| Dendritic | 262 | 0.382041 | -281.506 |

**Supplementary Table 2**

Statistics for PBMC data; for the differential entropy calculation, see **Methods**.

| Cell Type | Number of cells | % of total | Differential Entropy |
|---|---|---|---|
| Astrocyte | 54444 | 8.176518 | -285.773 |
| Endothelial Stalk | 39298 | 5.901859 | -271.857 |
| Endothelial Tip | 3818 | 0.573396 | -277.978 |
| Ependymal | 2157 | 0.323943 | -282.046 |
| Macrophage | 1695 | 0.254559 | -290.916 |
| Microglia | 4614 | 0.692941 | -275.472 |
| Mural | 12083 | 1.814651 | -270.937 |
| Neurogenesis | 2372 | 0.356232 | -257.468 |
| Neuron | 428051 | 64.28563 | -232.534 |
| Oligodendrocyte | 104773 | 15.73504 | -342.73 |
| Other (unlabeled) | 379 | 0.056919 | -281.542 |
| Polydendrocyte | 12174 | 1.828318 | -260.35 |

**Supplementary Table 3**

Statistics for adult mouse brain data; for the differential entropy calculation, see **Methods**.



**Preprint. Work in progress.**

| **Cell Type** | **Number of Cells** | **% of total** | **Differential Entropy** |
|---|---|---|---|
| Astrocyte | 34915 | 7.504067 | -293.16 |
| Ependymal | 2777 | 0.596844 | -274.99 |
| Immune/Blood | 14081 | 3.026343 | -289.20 |
| Neuron | 147059 | 31.60649 | -243.09 |
| Oligodendrocyte | 219220 | 47.11561 | -338.52 |
| Peripheral Glia | 16066 | 3.452967 | -328.23 |
| Vascular | 31163 | 6.697673 | -265.75 |

**Supplementary Table 4**

Statistics for developing and adolescent mouse CNS data; for the differential entropy calculation, see **Methods**.



**Preprint. Work in progress.**

| Cell Type | Uniform | *k*-means++ | SRS | Geometric sketching |
|---|---|---|---|---|
| Astrocyte | 1088 | 1090 | 389 | 1277 |
| Endothelial Stalk | 761 | 782 | 533 | 556 |
| Endothelial Tip | 84 | 107 | 175 | 815 |
| Ependymal | 33 | 68 | 102 | 165 |
| Macrophage | 43 | 31 | 47 | 262 |
| Microglia | 86 | 75 | 99 | 397 |
| Mural | 247 | 297 | 346 | 519 |
| Neurogenesis | 47 | 89 | 171 | 151 |
| Neuron | 8655 | 9746 | 10821 | 7975 |
| Oligodendrocyte | 2031 | 747 | 53 | 627 |
| Other (unlabeled) | 5 | 10 | 24 | 49 |
| Polydendrocyte | 237 | 275 | 557 | 524 |

**Supplementary Table 5**

Number of cells from each cell type in subsamples visualized in **Figure 2** from Saunders *et al*. (2018).

| Cell Type | Uniform | *k*-means++ | SRS | Geometric sketching |
|---|---|---|---|---|
| Astrocyte | 697 | 949 | 905 | 1194 |
| Ependymal | 49 | 115 | 339 | 614 |
| Immune/Blood | 265 | 418 | 350 | 466 |
| Neuron | 2982 | 4823 | 3911 | 4533 |
| Oligodendrocyte | 4371 | 1649 | 2273 | 1098 |
| Peripheral Glia | 332 | 322 | 259 | 230 |
| Vascular | 609 | 1029 | 1268 | 1170 |

**Supplementary Table 6**

Number of cells from each cell type in subsamples visualized in **Figure 2** from Zeisel *et al*. (2018).

[8] N. Halko, P.-G. Martinsson, and J. Tropp, “Finding Structure with Randomness: Probabilistic Algorithms for Constructing Approximate Matrix Decompositions,” *SIAM*

*Rev.*, vol. 53, no. 2, pp. 217–288, 2011.



**Preprint. Work in progress.**
